# Human airway organoids as a high-throughput screening platform for antiviral natural products discovery

**DOI:** 10.64898/2026.02.04.703746

**Authors:** Mathieu Hubert, Paola Haemmerli, Laurence Marcourt, Emy Lara-Quintero, Loan Arthaud, Luis-Manuel Quiros-Guerrero, Sarah Donnaray, Katia Rimensberger, Arnaud Gaudry, Elodie Alessandri-Gradt, Georgios Stroulios, Yves Cambet, François Prodon, Bohumil Maco, Samuel Constant, Salvatore Simmini, Antonio Grondin, Emerson Ferreira Queiroz, Sophie Clément, Jean-Luc Wolfender, Caroline Tapparel

## Abstract

Antiviral drug discovery for respiratory viruses is hindered by the lack of scalable physiologically relevant systems. Here, we report the first high-throughput screen of 764 natural plant extracts against respiratory syncytial virus (RSV) using human primary airway organoids as a relevant model. A parallel screen conducted in A549 cells allowed the identification of 70 extracts with organoid-specific antiviral activity from which 45 active phytocompounds were purified. We identified early- and late-acting antiviral compounds and demonstrated a polarization-dependent activity for some of them. Collectively, our results establish the use of airway organoids as a scalable first-line platform for high-throughput antiviral discovery and exploit the plant-derived chemical space as an underexplored source of RSV inhibitors.

## MAIN TEXT

Respiratory syncytial virus (RSV) is an enveloped negative-sense RNA virus belonging to the *Orthopneumovirus* genus in the *Pneumoviridae* family (*1*). Mainly transmitted via respiratory route (*2*), RSV causes essentially mild non-specific symptoms in healthy and immunocompetent adults (fever, cough, runny/stuffy nose, wheezing, dyspnea)(*3*) but can lead to severe lower respiratory tract infections (LRTIs) in at risk populations such as neonates and young children, elder adults (>60yo), immunocompromised patients and people living with chronic lung diseases (*3*). According to the World Health Organization (WHO), RSV infects 33 million children <5yo each year, causing 3.6 million associated hospitalizations with approximatively 100 000 RSV-attributable deaths (*4*). In this population, more than 20% of LRTIs are due to RSV (*4*), positioning this virus as a permanent public health concern. Substantial efforts have been made in RSV prevention. While two subunit vaccines (Abrysvo® and Arexvy®) and one mRNA-base vaccine (mRESVIA®) are available for prophylactic use in older adults and pregnant women (*5*), two new monoclonal antibodies (nirsevimab and clesrovimab) were recently FDA-approved to prevent RSV infection in neonates during their first RSV season (*6*). In contrast, therapeutic options remain extremely limited. Ribavirin was approved in 1986 to treat severe RSV infections (*7*), but its high cost and toxicity associated with a poor clinical efficacy have now restricted its use. To date, only the viral fusion protein (F) inhibitor ziresovir has successfully completed a phase 3 clinical trial in China (*8–10*), but resistance-associated mutations have already been reported (*9*). Thus, despite ongoing clinical development of other small-molecule candidates (*11*, *12*), effective therapeutic options remain extremely limited, underscoring the urgent need for novel antiviral candidates.

One important possible major limitation in the development of clinically effective antiviral therapies is that all previous antiviral RSV screenings were performed in immortalized cell lines that do not fully recapitulate the *in vivo* environment encountered by the virus (*13–18*). Because RSV primarily targets a structurally complex pseudostratified respiratory epithelium composed of pluripotent basal cells, ciliated cells and mucus-producing goblet cells, among other cell types (*19*), it is essential to perform screenings in physiologically relevant surrogate systems. Human airway epithelia differentiated at the air–liquid interface (ALI-HAE) constitute the gold standard *in vitro* model to study airborne viral infections (*20–22*) including RSV (*23*, *24*), but their cost, differentiation time and limited scalability preclude their use in high-throughput screenings. Another important limitation in the lack of treatment lies in the fact that available chemical libraries, despite their size, may not capture the full spectrum of chemical diversity found in nature and have therefore not been deeply explored for novel antiviral scaffolds. Natural products – particularly those derived from medicinal plants – represent a rich and still underexploited source of structurally diverse, biologically compatible molecules, and several plant extracts have already demonstrated antiviral activity across a range of viral families (*25–27*).

In this context, we show for the first time the feasibility of using apical-out airway organoids (AOAOs) (*28*) as a first instance physiologically relevant model to screen a chemically diverse library of several hundred metabolomically-profiled plant extracts for their antiviral activity against RSV under physiologically meaningful conditions. Part of this library has already been investigated for antiviral properties in our previous study (*30*),providing a valuable reference point for the present screening campaign.

### Apical-out airway organoids (AOAOs) as a relevant model to study RSV infection

We differentiated apical-out airway organoids (AOAOs) from human primary bronchial basal cells (BCs) following three steps (**Fig. 1A**): (i) the expansion of BCs in culture flasks, (ii) a 16-hour cell aggregation in AggreWell plates and (iii) a subsequent 15-day differentiation. Differentiation yielded homogeneous, spherical and well dispersed individual AOAOs (**Fig. S1A**) with a reproducible inter-batch average size of 70μm (**Fig. S1B** and **S1C**). We observed a lumen inside the organoids (**Fig. 1B**, top and dotted lines) and the honeycomb-like pattern of ZO-1 staining provided evidence for tight junction formation (**Fig. 1B**, bottom), indicating that BCs aggregation resulted in the generation of a polarized epithelium surrounding and enclosing an internal lumen. We confirmed the “apical-out” polarization by immunodetecting βIV-acetylated tubulin^+^ ciliated cells (green) and MUC5AC^+^ goblet cells (red) oriented toward the exterior (**Fig. 1C**). Scanning electron microscopy confirmed this externally oriented cilia layer interspersed with cilia-free goblet cells (**Fig. 1D**) and synchronized beating of motile cilia further demonstrated the functional integrity of the apical ciliary layer. To benchmark against the gold standard of the human airway epithelium cultured at air-liquid interface (ALI-HAE) *in vitro* models, we differentiated BCs into AOAOs and ALI-HAE in parallel and compared the expression of cell-specific markers for the three main epithelial cell types. Both ciliated (*Foxj1*, *Dnai1*) and goblet (*Muc5ac*, *Tff3*) cell-specific markers were significantly upregulated in AOAOs compared to non-differentiated BCs, concomitantly with the downregulation of the basal cell markers *Krt5* and *Tp63* (**Fig. 1E**). Notably, we observed similar differentiation efficiency between both models (**Fig. 1E**), validating AOAOs as a good surrogate for ALI-HAE. The susceptibility and permissiveness of AOAOs to RSV infection were assessed by exposing them to increasing doses of an RSV strain carrying a mCherry reporter gene (RSV-mCherry) for 2h at 37°C. After removal of the inoculum, we monitored mCherry fluorescence over time (**Fig. S1D** and **S1E**). RSV-exposed AOAOs displayed a dose-dependent mCherry signal (**Fig. 1F** and **S1F**) that peaked between 24 and 48h post-infection before declining (**Fig. 1G**). At the highest MOI, infection reached saturation at 24hpi with approximately 12% of infected cells (**Fig. S1G**), suggesting that innate immunity mechanisms are rapidly triggered to prevent viral spread into the whole organoid. Nevertheless, immune response is not sufficient to prevent cytopathic effects as demonstrated by the correlation between mCherry area and LDH release, a hallmark of cell death, at 24hpi (**Fig. S1H**). Kinetics of viral replication was further characterized by quantifying intracellular and extracellular viral RNA levels and confirmed the presence of a viral load peak as early as 24hpi (**Fig. 1H**), validating the mCherry signal previously observed as a surrogate of infection. Finally, a dose-dependent increase in the number of infectious particles was observed over time in the supernatant (**Fig. 1I**), demonstrating efficient production and release of a viral progeny in this model. Taken together, these findings prove that AOAOs constitute an efficient system to study RSV infection in a physiologically relevant context.

**Fig. 1.**
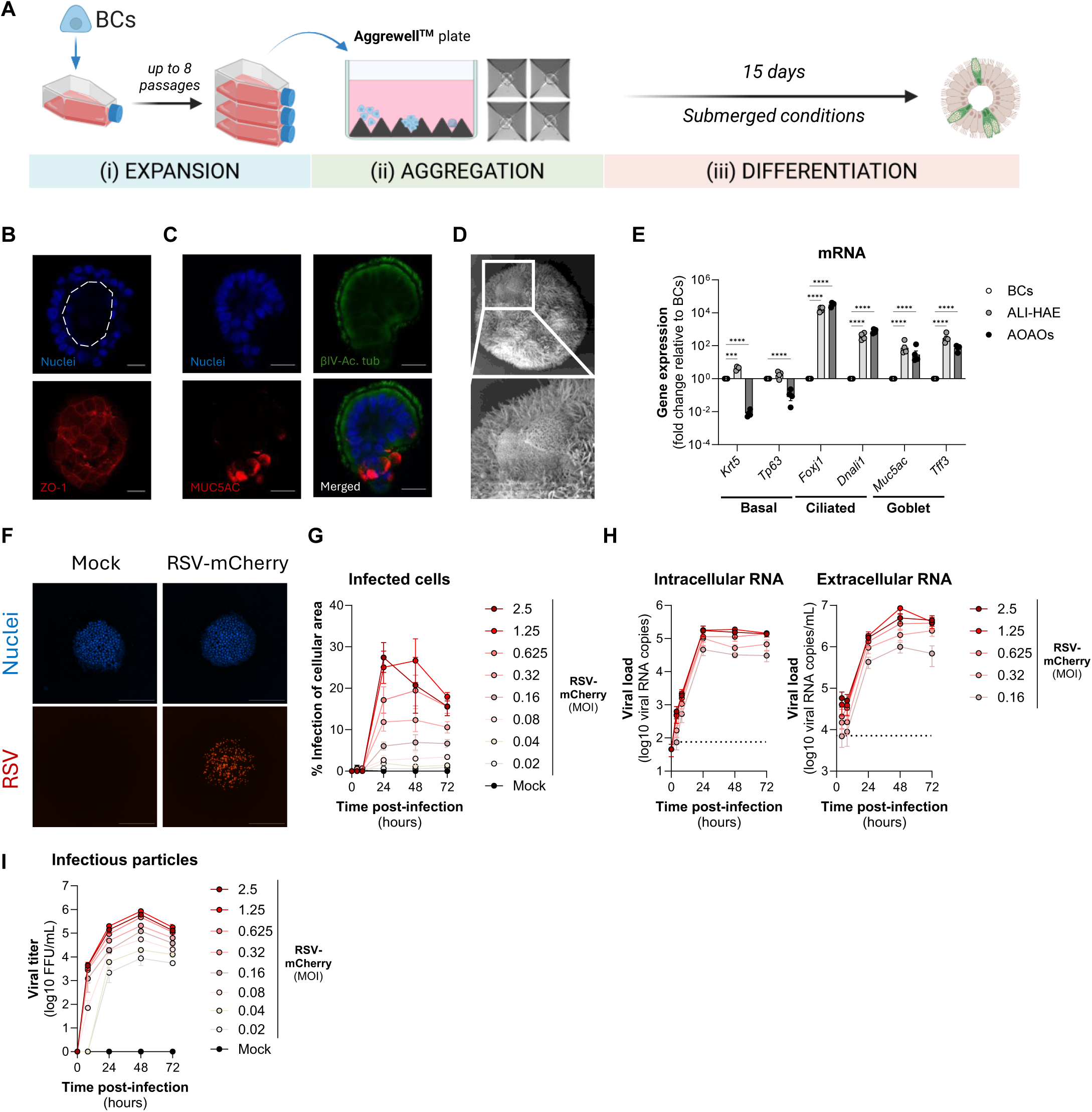
Characterization of apical-out airway organoids (AOAOs) and permissivity to RSV infection. **(A)** Schematic overview of AOAOs generation. Basal cells (BCs) were expanded in PneumaCult Ex-Plus medium for up to eight passages, aggregated for 16h in AggreWell™ plates, and differentiated for 15 days in Apical-Out Airway Organoid Medium. **(B)** Confocal micrographs of AOAOs after immunostaining of ZO-1 (red). Nuclei are stained in blue. Dotted line delineates the internal lumen. Micrographs were acquired at different Z to ensure the visualization of both the lumen and tight junctions. Scale bar: 20 μm. **(C)** Confocal micrographs of AOAOs after immunostaining of βIV-acetylated tubulin (green) and MUC5AC (red). Nuclei are stained in blue. Scale bar: 20 μm. **(D)** Scanning electron micrographs of fully differentiated AOAOs. Magnification: 500X (top) and 2000X (bottom). **(E)** Comparative gene expression analysis of cell-type-specific markers in AOAOs and ALI-HAE derived from the same BCs. Data were normalized to undifferentiated BCs and are presented as mean ± SEM from four independent experiments. Statistical test: two-way ANOVA with *** p=0.002; **** p<0.0001. **(F-I)** AOAOs were exposed to medium alone (Mock) or RSV-mCherry at indicated multiplicities of infection (MOI). At 4h, 8h, 24h, 48h and 72h post-infection, AOAOs and supernatants were collected. **(F)** Representative images of mock- or RSV-mCherry-infected (MOI 1.25) AOAOs at 24hpi. Nuclei are stained in blue. **(G)** Quantification of RSV infection by mCherry fluorescence measurement using a Cytation 5 microscope. Signal was normalized to the total cellular area (Hoechst). Results are expressed as mean±SEM of three independent experiments. **(H)** Quantification of intracellular and extracellular viral RNA copies from RSV-infected AOAOs. Results are expressed as mean±SEM of three independent experiments. **(I)** Viral titration of RSV-infected AOAOs’ supernatants onto A549 monolayers. Data are the result of one single experiment with two independent technical replicates.

### High-throughput screening of natural plant extracts against RSV in AOAOs

All previous RSV-addressed antiviral drug screens were performed in cell lines that do not recapitulate the microenvironment in which RSV replicates *in vivo* (*13–18*). Given both the sensitivity of AOAOs to RSV infection and the lack of clinically effective anti-RSV therapy, we aspired to discover new anti-RSV compounds by directly implementing a high-throughput screening (HTS) in this physiologically relevant model.

From a library of 1,600 well-documented and chemically characterized plant extracts originating from the Pierre Fabre library (*31*), we selected a subset of 764 chemically diverse samples (**Fig. SA**). We optimized experimental conditions to scale up RSV infections without inoculum removal in a HTS–compatible format (384-well plate) and identified a seeding density of 100 AOAOs per well (**Fig. 2A**) infected with a non-saturating multiplicity of infection of 0.05 (**Fig. 2B**) as optimal conditions to monitor RSV infection at a high-throughput scale in AOAOs. Thus, AOAOs were exposed to each natural plant extracts (50 μg/mL) in the presence of the optimal dose of RSV-mCherry (**Fig. 2C**) and fluorescence area was monitored 24h later, allowing us to identify extracts exhibiting both virus-directed and/or cell-directed antiviral activity. For comparison with conventional screening approaches, we conducted a parallel screen in A549 cells and included the vehicle alone and ribavirin (200 μM) in both models as internal negative and positive controls, respectively (*30*). Due to the low number of infected cells in organoids (**Fig. S1G**), variability issues prevented the calculation of any Z factor or Z-score for each plant extract. Nevertheless, we observed a significant and reproducible difference between negative and positive controls (**Fig. 2D**), enabling the implementation of a stringent ribavirin-matched threshold (<20% infection) to classify active extracts relative to vehicle-treated controls (**Fig. 2E**, pink area). Using these criteria, a total of 243 extracts (32%) were active in AOAOs (**Fig. 2E** and **2F**), while only 147 (19%) were active in A549 cells (**Fig. 2G-I**). Reproducibility of the hits was confirmed by significant correlation of the extract activity between two independent HTS in each model (**Fig. 2F** and **2I**), and comparison of reproducible hits in A549 and AOAOs revealed that 3% (23/764) and 9% (70/764) were specific to A549 and AOAOs, respectively, while 6% (44/764) were active in both systems (**Fig. 2J**). Notably, most active extracts (91.2%) exhibited no detectable toxicity (>75% viability) in AOAOs (**Fig. S3A**, top), while only 45% of active extracts met this criterion in A549 cells (**Fig. S3A**, bottom). These findings underscore the added value of performing primary HTS in physiologically relevant tissue culture models and highlight the limitations of cell line–based first-instance screens in which a substantial fraction of non-toxic and organoid-active extracts would likely go undetected. Most active extracts identified in AOAOs and A549 originated from green stems, leaves, roots, and barks (**Fig. S3B-D**).

**Fig. 2.**
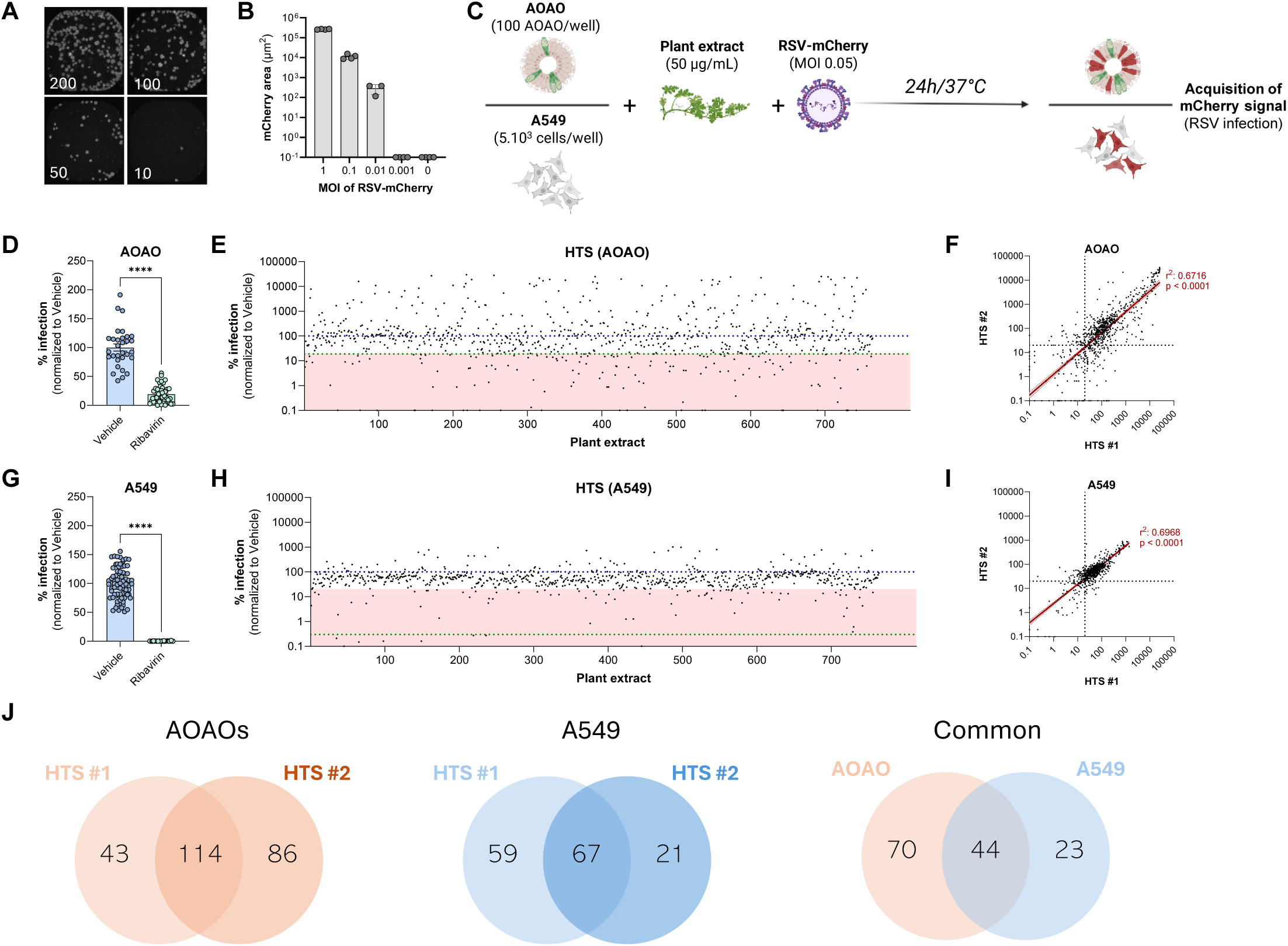
High-throughput screening of natural plant extracts against RSV in AOAOs. **(A)** AOAOs were seeded at different densities (200, 100, 50 and 10 AOAOs/well) in 384-well plates. Nuclei were stained with Hoechst and visualized using a Cytation 5 microscope. **(B)** AOAOs were exposed to indicated multiplicities of infection of RSV-mCherry. After 24h, mCherry signal was measured using Cytation 5 microscope and analyzed using Gen5 software. Results are expressed as mean±SD of 3-4 technical replicates of a single experiment. **(C)** Schematic workflow of the high-throughput screening of natural plant extracts against RSV in AOAOs. Briefly, 100 AOAOs/well were exposed to 50μg/mL of natural plant extracts in the presence of RSV-mCherry (MOI 0.05). Twenty-four hours later, mCherry area was measured as a surrogate of infection with Cytation 5 microscope and analyzed using Gen5 software. **(D)** Quantification of RSV-mCherry infection in vehicle- and ribavirin-treated AOAOs. Statistical test: non-parametric Mann-Whitney test with **** p<0.0001. **(E)** High-throughput screening in AOAOs. Each dot represents a plant extract. The green dotted line indicates the mean antiviral activity of ribavirin. The pink area represents the hits zone. Data are shown as mean of two independent experiments. **(F)** Reproducibility of the two independent HTS performed in AOAOs. Each dot represents a plant extract with the percentage of infection observed in the HTS#1 and HTS#2 in the x and y axes, respectively. Statistical test: simple linear regression. **(G)** Quantification of RSV-mCherry infection in vehicle- and ribavirin-treated A549 cells. Statistical test: non-parametric Mann-Whitney test with **** p<0.0001.**(H)** High-throughput screening in A549 cells. Each dot represents a plant extract. The green dotted line indicates the mean antiviral activity of ribavirin. The pink area represents the hits zone. Data are shown as mean of two independent experiments. **(I)** Reproducibility of the two independent HTS performed in A549 cells. Each dot represents a plant extract with the percentage of infection observed in the HTS#1 and HTS#2 in the x and y axes, respectively. Statistical test: simple linear regression. **(J)** Venn diagram of reproducible hits in the two independent HTS in AOAOs, A549 cells, and both models.

### Isolation and antiviral activity of plant-derived individual compounds

Plant extracts represent complex chemical matrices composed of hundreds of secondary metabolites (*32*). To identify individual compounds responsible for the antiviral activity of crude extracts against RSV, we selected four taxonomically and chemically distinct extracts with strong antiviral activity in both models (underground parts from *Ampelocissus arachnoida*, branches from *Clausena wallichii*), in A549 cells only (bark from *Litsea polyantha*), and in AOAOs only (leaves from *Vepris macrophylla*). From these extracts, we purified eight compounds from *Ampelocissus arachnoida* (A1–A8; **Fig. S4** and **S5**), twenty-one from *Clausena wallichii* (C1–C21; **Fig. S6** and **S7**), sixteen from *Litsea polyantha* (L1–L16; **Fig. S8** and **S9**), and six from *Vepris macrophylla* (V1–V6; **Fig. S10** and **S11**). All plant constituents were obtained at high purity and spanned diverse structural classes of NPs, including stilbene oligomers, coumarins, limonoids, carbazole and benzylisoquinoline alkaloids, phenylpropanoid derivatives, flavonoids, and furofuran lignans (**Table S1**). Their antiviral activity against RSV was assessed by dose-response assays in both AOAOs and A549 cells in parallel (**Fig. S12-S15** and **Table S2**) and led to the identification of a total of 15 compounds with reproducible activity in AOAOs, four of which were exclusively active in this model (**Fig. 3A**).

**Fig. 3.**
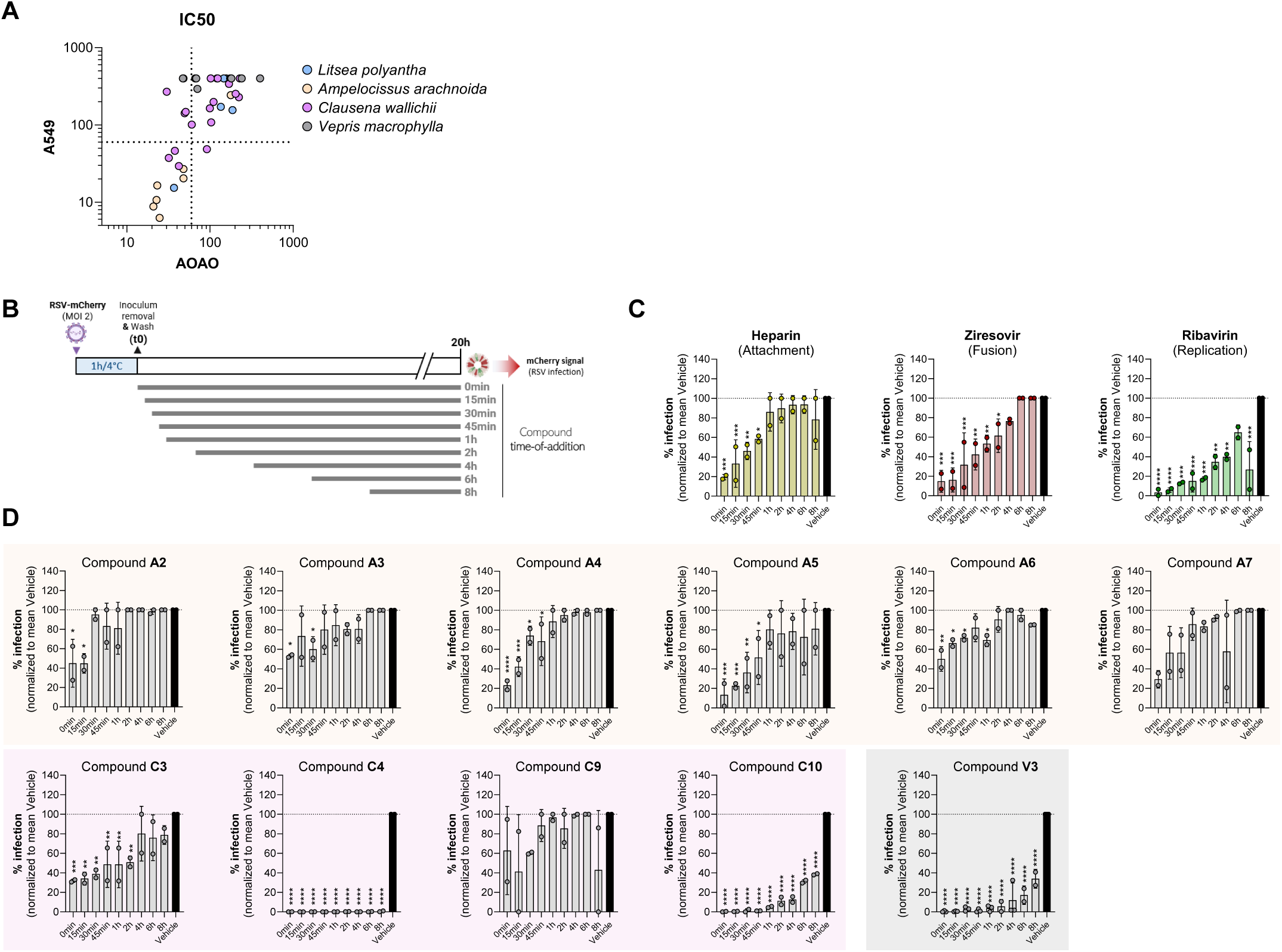
Antiviral activity of purified plant-derived compounds against RSV and kinetics of activity. **(A)** Dot plot representing the mean antiviral activity of each individual plant-derived compound based on IC_50_ values in AOAOs (x axis) and A549 cells (y axis). Dotted lines represent the 60μM threshold used to classify active compounds. Data are expressed as mean of two independent experiments. **(B)** Time-of-addition assay in AOAOs. Briefly, AOAOs (50 per well) were exposed to RSV-mCherry (MOI 2.5) for 1h at 4°C. After adsorption, inoculum removal and AOAOs washing, AOAOs were exposed to single-dose of selected plant-derived compounds at indicated time-points. After 20h, mCherry area was measured with Cytation 5 microscope and analyzed using Gen5 software. **(C)** Histograms showing the effects of heparin, ziresovir and ribavirin in the time-of-addition assay. Statistical test: one-way ANOVA with * p<0.0332; ** p<0.0021; *** p<0.0002; **** p<0.0001. Data are expressed as mean ± SD from two independent experiments. **(D)** Histograms showing the effects of compounds derived from *Ampelocissus arachnoida* (orange), *Clausena wallichi* (purple), and *Vepris macrophylla* (grey) in the time-of-addition assay. Statistical test: one-way ANOVA with * p<0.0332; ** p<0.0021; *** p<0.0002; **** p<0.0001. Data are expressed as mean ± SD from two independent experiments.

To identify the step of the RSV life cycle targeted by active compounds in AOAOs, we conducted a time-of-addition assay with 11 of the most active compounds in AOAOs (**Table S2**). Briefly, we exposed AOAOs to RSV-mCherry and added the maximal non-cytotoxic inhibitory concentration of each compound at indicated time-points (**Table S2** and **Fig. 3B**). At 20hpi, RSV infection was quantified via mCherry fluorescence. In the aim to delineate distinct stages of the RSV life cycle in AOAOs, we employed heparin, ziresovir, and ribavirin as reference inhibitors of viral attachment (*33*), fusion (*34*), and replication (*35*) respectively. As expected, heparin and ziresovir were mostly effective when applied early, within 45min and 2h post infection (hpi) respectively (**Fig. 3C**, left and middle panels), while ribavirin was still active when applied 8hpi (**Fig. 3C**, right panel). In contrast to dose–response conditions, two compounds (A7 and C9) lost their antiviral activity when added immediately after viral adsorption. Notably, all the other compounds isolated from *Ampelocissus arachnoida* acted early (15min to 1hpi) while those from *Clausena wallichii* and the single one from *Vepris macrophylla* displayed kinetics similar to ribavirin (**Fig. 3D**). This observation is consistent with the identification of structurally related compounds within each plant (**Table S1**) that inhibit RSV entry (stilbene oligomers from *Ampelocissus arachnoida*) or replication (carbazole alkaloids from *Clausena wallichii*) through shared mechanisms.

### Administration route

Contrary to organoids, air-liquid interface-cultures human airway epithelia (ALI-HAE) enable route-of-administration profiling by delivering compounds to the basolateral compartment, mimicking systemic (oral) exposure via the bloodstream, or to the apical surface, mimicking direct topical delivery to the airway lumen, as achieved by intranasal or inhaled administration. We selected A4 (hopeaphenol), A5 (α-viniferin), C3 (clausin K), C10 (3-formyl-2-hydroxy-7-methoxycarbazol) and V3 (phylligenin) as the five most promising antiviral compounds to investigate the polarization of their antiviral activity (**Fig. 4A**). After apical or basolateral treatment (**Fig. 4B**), antiviral efficacy was assessed by quantifying intra-and extracellular viral RNA levels by RT–qPCR and by measuring the infectivity of newly released viral particles. As toxicity in AOAOs only assess apical infection, we also checked the toxicity in ALI culture after compound administration from both sides. As expected, no cytotoxicity was detected upon apical administration, whereas basolateral treatment induced substantial toxicity for hopeaphenol (A4) or α-viniferin (A5) (∼40% loss of viability for both) and for clausine K (C3; ∼50%) at 72 h (**Fig. S3**). All compounds displayed robust antiviral activity when applied apically, consistent with direct exposure of the infected epithelium and in agreement with the AOAO-based screening results. In contrast, only α-viniferin (A5), 3-formyl-2-hydroxy-7-methoxycarbazol (C10) and phillygenin (V3) were also efficient when delivered basolaterally (**Fig. 4C**). The reference antivirals ribavirin and ziresovir were active *via* both apical and basolateral routes (**Fig. 4C**). Collectively, these results show that the antiviral activity of plant-derived compounds is mainly supported apically, opening perspectives of intra-nasal administration for antiviral treatment.

**Fig. 4.**
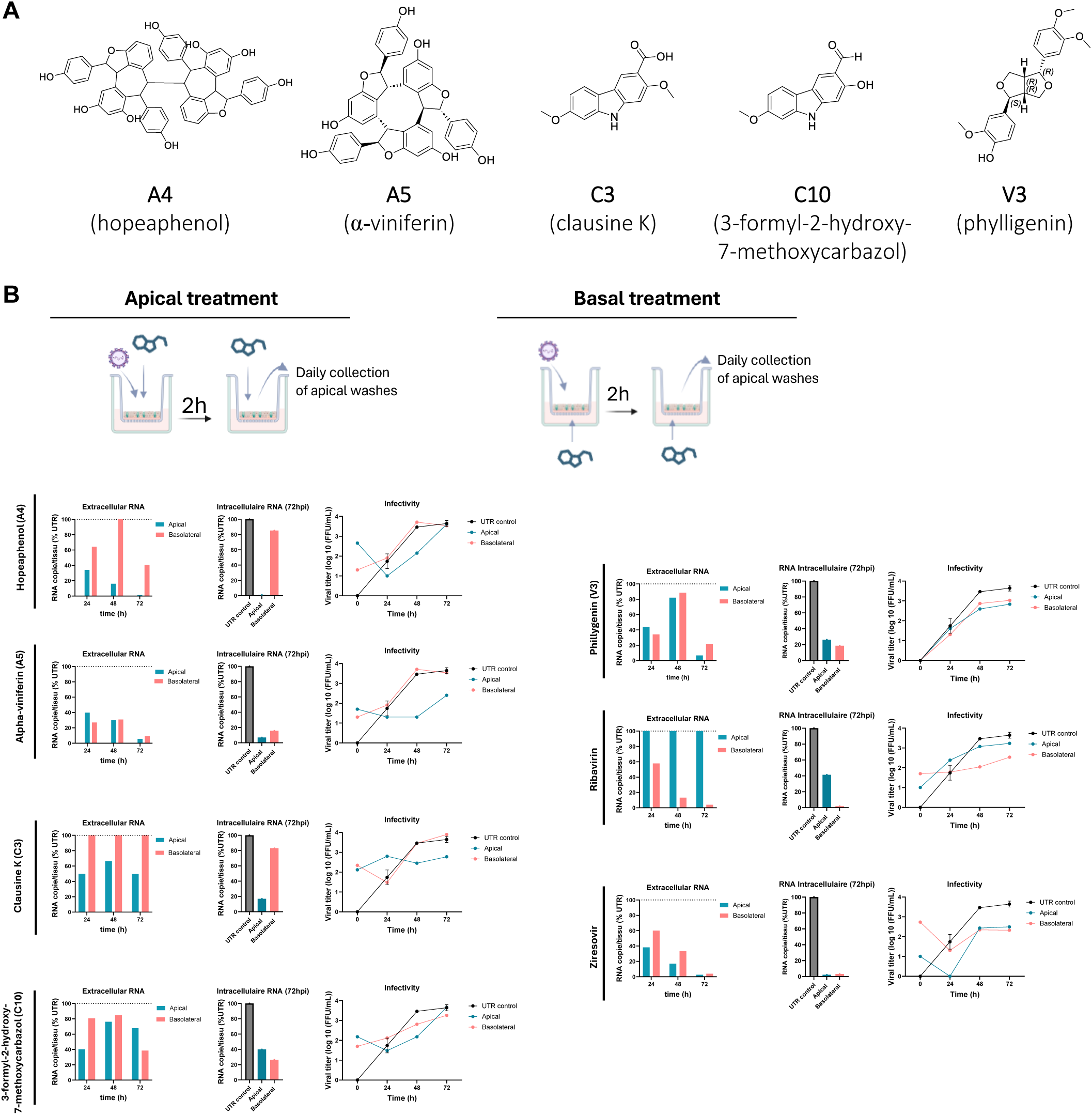
Polarization of plant-derived compounds’ antiviral activity in human airway epithelia differentiated at air-liquid interface (ALI-HAE). **(A)** Structure of the five selected compounds, namely hopeaphenol (A4), α-viniferin (A5), clausine K (C3), 3-formyl-2-hydroxy-7-methoxycarbazol (C10) and phylligenin (V3). **(B)** Schematic overview of the experimental workflow. Human airway epithelia cultured at air-liquid interface (ALI-HAE) were exposed to 5.10^3^ FFU for 2h at 37°C, washed, and daily treated with indicated compounds for 3 days via the apical or basolateral compartment. Each day, apical washes were collected for subsequent analysis. **(C)** Antiviral activity evaluated by extracellular viral RNA, infectious viral titers, and intracellular viral RNA levels at 78 h post-infection. Data represents one single experiment.

## DISCUSSION

In this study, we implemented apical-out airway organoids (AOAOs) as a scalable discovery tool for early-stage antiviral identification. Whereas in our previous work, AOAO were used to validate antiviral hits identified in A549 cells (*30*), they are established here as a primary high-throughput screening system. However, a side-by-side screening performed in a validated A549 assay (*30*) enabled direct comparison between both models.

We first validated the assay using ribavirin as a reference antiviral control, confirming the expected inhibition of RSV replication. Despite the intrinsic complexity of AOAO derived from primary bronchial cells (epithelial polarization, mucociliary defense, heterogeneous infection patterns), our AOAO-based assay exhibited a robust reproducibility across biological replicates when screening 764 plant extracts, comparable to that of the A549 assay. This performance shows that AOAO-based screening is not only biologically informative but also technically compatible with pharmacological discovery pipelines. Using stringent HTS thresholds (<20% of replication), 3% of extracts were active exclusively in A549 cells, 6% in both models, and 9% only in AOAOs. A549-restricted hits may reflect poorly translatable activities (e.g., chloroquine in SARS-CoV-2 (*36*)), whereas dual-active extracts represent high-confidence antiviral candidates. To our surprise, the largest fraction consisted of AOAO-specific hits, which would have been systematically missed in conventional antiviral screenings methods performed in cell lines. This observation suggests that epithelial differentiation state and tissue architecture could act as determinants of compound efficacy that are lacking in conventional 2D cellular assays. This conceptual advance is particularly illustrated by *Vepris macrophylla* and its major constituent phillygenin (V3), a furanolignan whose antiviral activity was detected exclusively in AOAOs but not in A549 cells. This finding is consistent with previous *in silico* predictions of anti-RSV activity and with its lack of efficacy in another cell line (Hep-G2) in which studies reported either no efficacy or only weak antiviral activity (*37*, *38*). Independent *in vivo* evidence further supports the relevance of phylligenin to be uses as an antiviral treatment since its combination with baicalin improved respiratory outcomes in a chicken infectious bronchitis virus model (*39*). Importantly, phillygenin also shows a narrow structure–activity relationship, being the only active furanolignan despite close similarity to related analogues (V4–V6). Collectively, these findings establish phillygenin as a proof of concept for the AOAO-based assay, highlighting the ability of physiologically relevant models to reveal antivirals missed by conventional screening approaches.

Beyond AOAO-specific hits, we identified other antivirals from diverse chemical families, notably stilbene derivatives. These stilbene derivatives were active in both AOAOs and A549 cells and were shown to inhibit multiple enveloped viruses, supporting their potential as broad-spectrum antivirals (*40*, *41*). We also identified isoquinoline alkaloids, including reticuline, whose antiviral activity has not previously been reported but which belongs to the same family than berberine, a known RSV inhibitor (*42*). In addition, carbazole alkaloids exhibited strong antiviral activity against RSV in AOAOs, further expanding the chemical space of RSV inhibitors. The bioactivity profiles of these fully characterized constituents also confirm that all the selected plant extracts exhibit a level of chemical diversity that is well suited to the discovery of anti-RSV molecules.

Importantly, our AOAO-based assay enabled not only compound prioritization but also early mechanistic profiling, revealing similar modes of action within chemical families. Especially, stilbene oligomers consistently acted at early stages of the viral replication cycle. Given their molecular size, this profile could be compatible with effects on viral particles at early entry steps rather than intracellular replication (likely reflecting limited cellular penetration). This interpretation is supported by our previous work showing that trans-δ-viniferin derivatives exert direct virucidal activity against influenza A virus (H1N1) by disrupting viral envelope (*40*). While our study provides initial mechanistic insights, no antiviral mechanism has yet been experimentally validated for these compounds in the context of RSV, and precise molecular targets remain to be defined. Finally, all hits identified in AOAO were also active in *ex vivo* human airway epithelium following apical administration and showed no detectable toxicity in either system. Importantly, the AOAO-based screening also allowed identification of active candidates when administered basally in ALI models, as illustrated by phillygenin (V3). While immune and stromal components also contribute to RSV pathogenesis *in vivo*, an epithelial-focused model provides a biologically relevant system for primary antiviral screening. Beyond RSV, this assay may be broadly applicable to the discovery of antivirals against other respiratory viruses that critically depend on epithelial differentiation state and tissue architecture, as well as other respiratory pathogens.

## MATERIALS AND METHODS

### Cells and viruses

A549 cells (kind gift from Pr. Mirko Schmolke, Geneva, Switzerland) were grown in Dulbecco’s Modified Eagle Medium (DMEM, Gibco) supplemented with 10% fetal bovine serum (FBS, Gibco), and 1% penicillin/streptomycin (Gibco) at 37°C; 5% CO_2_. Human primary bronchial airway basal cells (BCs) were collected from an adult healthy donor (#AB0839, Epithelix) and expanded in PneumaCult Ex-Plus Medium (STEMCELL Technologies) at 37°C; 5% CO_2_. RSV-mCherry (kind gift from Pr. J.-F. Elouët, Institut National de la Recherche Agronomique, Jouy-en-Josas, France)(*43*) was propagated in A549 cells in DMEM supplemented with 2.5% FBS and 1% penicillin/streptomycin at 37°C; 5% CO_2_. For viral stocks preparation, A549 cells were scraped and cell debris-containing supernatants were vortexed for 5min. After clarification, viral stocks were cryopreserved at −80°C in the presence of cryoprotective agents. Viral titers were determined by focus-forming assay as previously described (*30*) and expressed as foci-forming units per mL (FFU/mL).

### Apical-out airway organoids (AOAOs)

After three to eight passages in PneumaCult Ex-Plus medium, 7.10^5^ basal cells (BCs) were seeded in 6-well AggreWell 400 plates (STEMCELL Technologies) in Apical-Out Airway Organoid Medium (AOAO-M, STEMCELL Technologies) and centrifuged at 100g for 5min. Before cell seeding, AggreWell plates were pre-treated with Anti-Adherence Rinsing solution (STEMCELL Technologies) to prevent adherence of BCs. Cell aggregates were incubated for 15 days at 37°C; 5% CO_2_ with medium renewal every two days. When fully differentiated, AOAOs were gently collected, filtered through a 37 µm cell strainer (STEMCELL Technologies) and resuspended in AOAO-M for manual counting and further experiments.

### Air-liquid interface human airway epithelia (ALI-HAE)

After three to eight passages in PneumaCult Ex-Plus medium, 4×10^4^ basal cells (BCs) were seeded onto 0.4 μm polyester Transwell® inserts (#3470, Corning) in PneumaCult Ex-Plus medium. When confluency was reached, basal medium was replaced with PneumaCult ALI medium (ALI-M; STEMCELL Technologies) and apical medium was removed for further differentiation at air-liquid interface (ALI). After 21 days, fully differentiated ALI human airway epithelia (ALI-HAE) were used for further experiments.

### Viral infections

All infections performed in AOAOs were conducted in AOAO-M without heparin to prevent virus inhibition. To assess organoid permissiveness, AOAOs were incubated with RSV-mCherry at indicated multiplicities of infection (MOI) for 2h at 37°C, washed with Dulbecco’s Phosphate Buffer Saline without calcium and magnesium chloride (D-PBS; Sigma) and resuspended in AOAO-M. For cell imaging, AOAO nuclei were stained with Hoechst 33342 (Invitrogen). At 4h, 8h, 24h, 48h and 72h post-infection (hpi), AOAOs were collected for cell imaging and measurement of both Hoechst and mCherry fluorescence (Biotek Cytation 5 Cell imaging multimode reader, Agilent), immunofluorescence, scanning electron microscopy, qRT-PCR, flow cytometry, and supernatants were collected for qRT-PCR and viral titration (see respective sections below).

### Antiviral activity assessment

For dose-responses of plant-derived compounds, AOAOs were exposed to RSV-mCherry (MOI 0.32) in the presence of vehicle alone or 2-fold serial dilutions of individual compounds. After 20h, RSV-mCherry^+^ area was measured and normalized to the total Hoechst^+^ cell area to calculate a percentage of infection. In A549 cells, dose-responses were assessed following the experimental procedures described previously (*30*). For time-of-addition assays, RSV infections were synchronized by exposing AOAOs to RSV-mCherry (MOI 2) for 1h at 4°C followed by PBS washing. The maximal inhibitory but non-cytotoxic dose of each individual plant-derived compound (**Table S2**) was added immediately or after 15min, 30min, 45min, 1h, 2h, 4h, 6h, 8h. After 20h, the percentage of infection was calculated as above. For ALI-HAE infections, tissues were apically exposed to 5×10^3^ FFU for 2h at 37 °C in ALI-M without heparin. After adsorption, ALI-HAE were washed and subsequently treated with plant-derived compounds at indicated doses either apically or basolaterally. Daily, apical washes were collected, and treatments were refreshed. Viral replication was monitored by measuring extracellular viral RNA and infectious RSV particles (see respective sections below). At 78 h post-infection, tissues were lysed for quantification of intracellular viral RNA levels (see respective sections below).

### Immunofluorescence and optical microscopy

For intracellular staining, AOAOs were fixed in 4% paraformaldehyde (PFA; Reactolab), permeabilized with PBS containing 1% Triton (AppliChem) and blocked in PBS with 5% BSA (AppliChem), each step performed for 3h at RT. For primary staining, AOAOs were incubated overnight at 4°C with rabbit anti-acetylated βIV-tubulin (ab179504, abcam) and mouse anti-MUC5AC (ab3649, abcam) antibodies diluted in PBS supplemented with 0.1% BSA, 0.2% Triton and 0.1% Tween20 (AppliChem). After washing, samples were incubated for 3h at RT in the dark with Alexa Fluor 488-conjugated goat anti-rabbit (A11008, Invitrogen) and Alexa Fluor 555-conjugated goat anti-mouse (A31555, Invitrogen) secondary antibodies. Nuclei were stained with Hoechst 33342. Following washes, AOAOs were mounted on slides using Prolong Gold Antifade mounting medium (Invitrogen). Fluorescently labeled samples were acquired on a Leica Stellaris 5 upright confocal laser scanning microscope. Live imaging was performed using an inverted Zeiss Axio Observer Z1. Images were analyzed using Fiji software (Java 1.6.0_24).

### Scanning electron microscopy

Organoids were fixed in 2.5% glutaraldehyde and 2.0% paraformaldehyde mixture in 0.1M sodium cacodylate buffer pH 7.4 for 1 h at room temperature. After 3 washes with 0.1M sodium cacodylate buffer pH 7.4 samples were incubated with 1% tannic acid in 0.1M sodium cacodylate buffer pH 7.4 for 1 h at RT, washed 3 times with 0.1M sodium cacodylate buffer pH 7.4 and post-fixed with 1% osmium tetroxide in 0.1M sodium cacodylate buffer pH 7.4 for 1 h at RT. After 3 washes with 0.1M sodium cacodylate buffer pH 7.4 samples were dehydrated with graded series of ethanol: 50% for 10 min, 70% for 10 min, 90% for 10 min and 100% ethanol 3 times 10 min. Organoid samples were immediately transferred into critical-point dryer (Quorum K850 CPD). Fully dehydrated organoids were deposited on double sided adhesive carbon tape attached to the SEM flat aluminum stubs and immediately coated with 2-3 nm layer of gold and imaged with Helios 660 Nanolab DualBeam SEM (ThermoFisher Scientific) at 2-5 kV of accelerator voltage and 0.1 nA current using ETD secondary electron detector and pixel dwell time of 5 µsec.

### RT-qPCR

Samples were lysed in TRK lysis buffer (E.Z.N.A Viral RNA Extraction Kit; Omega Bio-Tekand) and total RNA was extracted according to the manufacturer’s recommendations. Gene expression was quantified by TaqMan-based one-step reverse transcription quantitative PCR (RT-qPCR; QuantiTect RT, Qiagen) using specific primers and probes for basal cell markers (*Krt5*, Hs00361185_m1; *Tp63*, Hs00978343_m1), ciliated cell markers (*Foxj1*, Hs00230964_m1; *Dnali1*, Hs00185750_m1), and goblet cell markers (*Muc5AC*, Hs01365616_m1; *Tff3*, Hs00902278_m1), with *Gapdh* (#4310884E) used as a housekeeping gene (all from Thermo Fisher Scientific). RSV viral loads were quantified using N gene–specific primers (forward: 5′-TGC TAA GAC YCC CCA CCG TAA C-3′; reverse: 5′-GGA TTT TTG CAG GAT TGT TTA TGA-3′) and a Yakima Yellow–labeled probe (5′-CAC TTG CCC TGC WCC A-BHQ-3′). Amplification was performed with the following cycling conditions: 10 min at 95°C, followed by 45 cycles of 20 s at 60°C and 30 s at 72°C (7500 Fast Real-time PCR system, Applied Biosystems). Expression levels were expressed using the comparative ΔΔCt methodology, using BCs as the reference condition.

### Flow cytometry

AOAOs were treated with 0.05% trypsin-EDTA (Gibco) at 37°C until complete dissociation. Single cells suspensions were fixed with PFA 4% for 20min at RT and resuspended in PBS. Samples were acquired on a Cytoflex instrument (Beckman Coulter) and analyzed based on their mCherry fluorescence. Data were analyzed using FlowJo software (v10.10.0) and expressed as a percentage of mCherry^+^ infected cells.

### Viral titration

To quantify infectious RSV particles produced by AOAOs and ALI-HAE, culture supernatants were collected and immediately applied onto confluent A549 monolayers. As references, serial dilutions of a viral stock of known titer (expressed in FFU/mL) were applied under identical conditions. After 24h, mCherry fluorescence was measured in sample wells and interpolated against the standard curve to deduce equivalent titers expressed as FFU/mL.

### Plant material

The plant used in this study originates from the Pierre Fabre Library (PFL)(Pierre Fabre Laboratories, Toulouse, France) as a large and curated library established between 1998 and 2015 for natural product–based drug discovery and comprising more than 17,000 unique samples (*31*). Plant materials were air-dried, ground, and stored under controlled conditions in a secure facility. Metadata, including geographical origin, sample identifiers, and quantities, are maintained in the PFL internal database, and reference plant material is conserved to enable retrospective botanical verification. The historical collection complies with the Nagoya Protocol and is registered with the European Commission (accession no. 03-FR-2020)(*44*). Plant materials from this collection are available in several formats, including dried natural extracts, corresponding dried plant material, and ready-to-use extract solutions solubilized in dimethyl sulfoxide (DMSO; original extracts), as previously reported. In the present study, screening experiments were performed using the DMSO-solubilized natural extracts.

### Extracts from the Pierre-Fabre Library

Original ethyl acetate (EtOAc) natural extracts (NEs) from Pierre Fabre Laboratories used for the screening in A549 and AOAO were generated from dried plant material using a previously described standardized extraction protocol (*31*). This protocol was specifically designed and already optimized to minimize the presence of known interfering compounds, including condensed tannins. Extracts were dissolved in DMSO at a final concentration of 5 mg.mL⁻¹ and transferred into 96-well plates (Corning). In this study, a total of 764 plant extracts were evaluated by high-throughput screening, covering 381 species distributed across 275 genera and 102 botanical families. All were part of a set of 1,600 extracts already subjected to systematic metabolite profiling by UHPLC–PDA–HRMS/MS (*31*).

### High-throughput screening of the natural extracts

In 384-well plates, 100 AOAO per well were seeded in 384-well plate (Corning) and exposed to RSV-mCherry (MOI 0.05) in the presence of 50µg/mL of each natural plant extract or 1% dimethylsulfoxyde (DMSO, Sigma) or ribavirin (*30*) as negative and positive controls, respectively. Hoechst was added in the medium, permitting the detection of organoids. After 20h, Hoechst and mCherry areas were directly measured in the 384-well format (Cytation 5) and the percentage of infection was calculated as above. The parallel HTS in A549 cells was assessed following the experimental procedure we described previously (*30*).

### Scale-up extraction of the four selected plants

To enable phytochemical investigation and compound isolation, the four selected plant species *Clausena wallichii* (branches), *Ampelocissus arachnoida* (undergroundparts), *Litsea polyantha* (bark) and *Vepris macrophylla* (leaves) were subjected to scaled-up extraction procedures. Extractions were performed using an Accelerated Solvent Extraction (ASE) system equipped with a 100 mL stainless steel pressure-resistant extraction cell, according to a procedure we previously described (*30*). Briefly, dried and ground plant materials were sequentially extracted with solvents of increasing polarity: HPLC-grade hexane, ethyl acetate (EtOAc), and methanol (MeOH)(Fisher Chemicals, Reinach, Switzerland). Resulting extracts were concentrated under reduced pressure at 40 °C using a rotary evaporator. For *Ampelocissus arachnoida*, 35g of green stems yielded 1g of hexane extract, 10g of EtOAc extract, and 10g of MeOH extract. For *Clausena wallichii*, 24.6g of dried root material yielded 91mg of hexane extract, 551mg of EtOAc extract, and 1.31g of MeOH extract. For *Litsea polyantha,* 47g of dried root material yielded 88mg of hexane extract, 338mg of EtOAc extract, and 400mg of MeOH extract. For *Vepris macrophylla*, 27g of dried root material yielded 652mg of hexane extract, 678mg of EtOAc extract, and 4.3g of MeOH extract.

### Metabolite profiling of the four selected plants

The ethyl acetate extracts obtained from the scaled-up extraction of four selected plant species (*Clausena wallichii*, *Ampelocissus arachnoida*, *Litsea polyantha* and *Vepris macrophylla*) were analyzed using an optimized UHPLC–PDA–CAD–HRMS/MS workflow that we recently described (*30*) to assess their metabolite profiles and subsequent comparison with original extracts (*31*) to confirm their chemical composition. To isolate and purify plant-derived compounds, two different strategies were applied depending on the chemical composition complexity of the extracts (see respective sections below). For the less complex extracts (*Litsea polyantha* and *Vepris macrophylla*), compound isolation was achieved using a one-step strategy. For the more complex extracts (*Clausena wallichii* and *Ampelocissus arachnoida*), an additional separation step was required prior to isolation.

### One-step isolation of plant-derived compounds of *Litsea polyantha* and *Vepris macrophylla*

The main constituents of *Litsea polyantha* and *Vepris macrophylla* were isolated from ethyl acetate extracts. Extracts of *Litsea polyantha* (3 mg mL⁻¹ in HPLC-grade methanol) and *Vepris macrophylla* (3 mg mL⁻¹ in HPLC-grade methanol) were analyzed using a UHPLC system equipped with a reversed-phase C18 column (Waters ®, 150 × 2.1 mm i.d., 1.7 µm). The flow rate was set to 0.4 mL min⁻¹, and a binary solvent system consisting of water with 0.1% formic acid (A) and acetonitrile with 0.1% formic acid (B) was employed (*45*). An initial linear gradient of solvent B was applied to both extracts as follows [t (min), %B]: 0.00, 5; 0.20, 5; 8.00, 100; 9.00, 100; 9.50, 5; 10.00, 5. For *Litsea polyantha*, the method was subsequently optimized to improve compound separation using the following gradient program [t (min), %B]: 0.00, 5; 0.20, 5; 0.50, 10; 8.00, 20; 9.00, 100; 9.50, 100; 9.60, 5; 10.00, 5. For *Vepris macrophylla*, chromatographic separation was optimized using the following gradient program [t (min), %B]: 0.00, 5; 0.20, 5; 0.50, 30; 9.00, 45; 9.10, 100; 9.50, 100; 9.60, 5; 10.00, 5. Based on the optimized analytical conditions, gradient transfers were calculated to enable semi-preparative chromatographic separation of both extracts (*46*). The system was equipped with a guard column (Waters® C18, 10 × 19 mm), a C18 analytical column (Waters® XBridge, 250 × 19 mm i.d., 5 µm). The flow rate was set to 17 mL min⁻¹, with an average run time of approximately 60 min. The mobile phase consisted of H₂O containing 0.1% formic acid (A) and ACN containing 0.1% formic acid (B). For *Litsea polyantha*, separation was performed in three injections (3 x 50.0 mg) and semi-preparative chromatographic separation afforded 1.2 mg of compound **L1** (RT 17.5 min), 1.1 mg of **L2** (RT 18 min), 0.9 mg of **L3** (RT 18.3 min), 0.3 mg of **L4** (RT 20 min), 0.3 mg of **L5** (RT 21 min), 0.2 mg of **L6** (RT 22.0 min), 0.8 mg of **L7** (RT 33 min), 0.3 mg of **L8** (RT 34 min), 0.1 mg of **L9** (RT 35 min), and 0.4 mg of **L10** (RT 40.0 min). For *Vepris macrophylla*, separation was performed in two injections (2 x 50 mg) with a dry load injection using a dry-load injection (*47*), and fractions were collected every 12 mL. A similar procedure than for *Litsea. polyantha* yielded 0.9 mg of compound **V1** (RT 19.5 min), 1.5 mg of compound **V2** (RT 19.5 min), 13.0 mg of compound **V3** (RT 26 min), 3.8 mg of compound **V4** (RT 39 min), 0.7 mg of compound **V5** (RT 41 min), and 0.1 mg of compound **V6** (RT 41.4 min).

### Multistep isolation of plant-derived compounds of *Clausena walichii* and *Ampelocissus arachnoida*

The main constituents of *Clausena wallichii* and *Ampelocissus arachnoida* were isolated from ethyl acetate extracts. Extracts of *Clausena wallichii* (3 mg mL⁻¹ in HPLC-grade methanol) and *Ampelocissus arachnoida* (3 mg mL⁻¹ in HPLC-grade methanol), chromatographic analyses were performed using the same HPLC system (*45*). For *Clausena wallichii*, separations were carried out on an C18-HP column (Interchim®, 250 × 4.6 mm i.d., 15 µm), at a flow rate of 2 mL min⁻¹, using a binary solvent system consisting of 0.1% formic acid (FA) in water (A) and 0.1% FA in HPLC-grade methanol (B). An initial linear gradient of solvent B was applied as follows [t (min), %B]: 0.00, 2; 2.00, 2; 7.00, 2; 50.00, 98; 55.00, 98, followed by a re-equilibration step (60.00, 2). This method was subsequently optimized to improve compound separation using the following gradient program [t (min), %B]: 0.00, 2; 2.00, 2; 7.00, 55; 45.00, 98; 50.00, 98; 55.00, 98, followed by re-equilibration steps (57.00, 2; 60.00, 2). Based on the optimized analytical conditions, a gradient transfer was calculated to enable flash chromatographic separation of the *Clausena wallichii* extract (*46*).

The *Clausena wallichii* extract (0.508 g) was separated on a Puriflash C18-HP column (Interchim®, 200 × 30 mm, i.d.,15 µm), using an Interchim system. The flow rate was set to 60 mL min⁻¹, and a binary solvent system consisting of 0.1% aqueous FA [A] and 0.1% FA in HPLC-grade MeOH [B] was used. A gradient (v/v) of solvent B was applied as follows [t (min), %B]: 0.00, 2; 1.00, 2; 6.00, 55; 43.00, 98; 53.00, 98; 55.00, 2; 60.00, 2. Fractions collection was performed using a fixed fraction volume of 20 mL per tube. After combining tubes based on UV traces at 254 nm and 280 nm, a total of 35 fractions was obtained (fraction CWE-F1 to CWE-F35). This separation yielded 19.3 mg of **C3** (RT 17.0 min), 14.7 mg of **C10** (RT 21.1 min), 8.7 mg of **C18** (RT 26.5 min), 4.7 mg of **C17** (RT 27.3 min), and 11.5 mg of **C21** (RT 35.2 min). The fraction collected at RT 24.4 min (CWE-F27, 3.7 mg) was separated using a Shimadzu semi-preparative system using dry load injection on an X-Bridge C18 column (250 × 10 mm i.d., 5 µm) equipped with a Waters C18 guard cartridge holder (5 × 10 mm i.d., 5 µm); the solvent system consisted of HPLC-grade MeOH (B) and water (A), both containing 0.1% FA, using a linear gradient of 40–45% MeOH, to afford **C3** (0.6 mg, RT 14.45 min), **C13** (0.6 mg, RT 29.0 min), **C15** (0.6 mg, RT 32.3 min), and **C16** (1.8 mg, RT 35.00 min), respectively. Subsequent separations were performed using the same solvent system and column, but with different linear gradient compositions. Fraction CWE-F10 (12.4 mg, RT 19.1 min) was separated using the same linear gradient as above to yield **C3** (1.7 mg, RT 14.15 min), **C4** (1.9 mg, RT 15.2 min), and **C7** (0.6 mg, RT 19.45 min). Fraction CWE-F13 (11.3 mg, RT 22.2 min) was separated using a linear MeOH gradient of 35–40% to yield **C3** (0.5 mg, RT 17.3 min), **C8** (1.2 mg, RT 24.4 min), **C9** (1.5 mg, RT 28.2 min), **C11** (0.6 mg, RT 30.0 min), and **C12** (0.5 mg, RT 30.45 min). The following fractions were all separated using a linear MeOH gradient of 45–50%. Fraction CWE-F26 (3.5 mg, RT 23.5 min) was separated to yield **C3** (0.3 mg, RT 12.3 min), **C9** (0.2 mg, RT 16.45 min), **C13** (2.1 mg, RT 21.30 min), **C14** (0.6 mg, RT 22.3 min), and **C16** (0.7 mg, RT 26.0 min). Fraction CWE-F29 (10.8 mg, RT 28.0 min) was separated to yield **C17** (3.1 mg, RT 30.0 min) and **C19** (2.8 mg, RT 33.4 min). Fraction CWE-F30 (2.7 mg, RT 28.3 min) was separated to yield **C17** (0.4 mg, RT 31.1 min). Fraction CWE-F33 (14.4 mg, RT 34.4 min) was separated using an isocratic mode at 55% MeOH for 75 min to yield **C21** (5.3 mg, RT 36.1 min) and **C20** (2.4 mg, RT 38.15 min).

The following fractions were all separated using a linear MeOH gradient of 30–35%. Fraction CWE-F02 (1.6 mg, RT 13.1 min) was separated to yield **C5** (0.6 mg, RT 35.3 min). Fraction CWE-F05 (2.7 mg, RT 14.2 min) was separated to yield **C1** (0.4 mg, RT 17.5 min), **C2** (0.3 mg, RT 23.0 min), and **C6** (0.5 mg, RT 35.4 min).

For *Ampelocissus arachnoida* extract, HPLC separations were performed on a phenyl column (XBridge®, 250 × 4.6 mm i.d., 5 µm) equipped with a phenyl guard cartridge (10 × 4.6 mm i.d., 5 µm) at a flow rate of 1 mL min⁻¹, using a binary solvent system consisting of 0.1% FA in water (A) and 0.1% FA in HPLC-grade acetonitrile (B). An initial linear gradient of solvent B was applied as follows [t (min), %B]: 0.00, 2; 2.00, 2; 7.00, 25; 45.00, 40; 50.00, 98; 55.00, 98, followed by a re-equilibration step (57.00, 2; 60.00, 2). The method was then optimized using the following gradient program [t (min), %B]: 0.00, 2; 2.00, 2; 7.00, 55; 45.00, 98; 50.00, 98; 55.00, 98, followed by re-equilibration steps (57.00, 2; 60.00, 2). A gradient transfer was subsequently calculated to allow semi-preparative HPLC chromatographic separation of the *Ampelocissus arachnoida* extract. The *Ampelocissus arachnoida* extract (0.1044 g) was separated on a Kinetex Phenyl column (Shimadzu®, 250 × 21 mm i.d., 5 µm) using semi-preparative HPLC using a preparative LC system (LC-20 AP, Shimadzu, Kyoto, Japan)(*45*). The flow rate was set to 19 mL min⁻¹ and a binary solvent system consisting of 0.1% aqueous FA [A] and 0.1% FA in HPLC-grade ACN [B] was used. A gradient (v/v) of solvent B was applied as follows [t (min), %B]: 0.01, 2; 2.72, 2; 7.66, 25; 45.17, 40; 50.11, 98; 55.04, 98; 57.02, 2; 60.00, 2. Fraction collection was performed using a fixed fraction volume of 19 mL per tube. After combining tubes based on UV traces at 254 nm and 280 nm, a total of 21 fractions were obtained. This separation yielded 0.8 mg of **A1** (RT 22.25 min), 0.5 mg of **A2** (RT 27.0 min), 1.0 mg of **A3** (RT 28.0 min), 5.9 mg of **A4** (RT 35.0 min), and 9.3 mg of **A5** (RT 45.5 min).

The *Ampelocissus arachnoida* fraction collected at RT 31.3 min (AAE-F12, 6.3 mg) was separated at the semi-preparative scale using an HPLC 1260-II system on an BEH Phenyl OBD Prep Column (XBridge®, 250 × 10 mm i.d., 5 µm) equipped with a phenyl guard cartridge holder (5 × 10 mm i.d., 5 µm), using a solvent system composed of HPLC-grade ACN (B) and H₂O (A), both containing 0.1% FA, under isocratic conditions at 25% ACN, to yield **A6** (2.1 mg, RT 28.3 min). Subsequent separations were performed using the same solvent system and column, but with different linear gradient compositions. Fraction AAE-F15 (4.3 mg, RT 36.3 min) was separated using an isocratic mode at 30% ACN to yield **A7** (2.6 mg, RT 21.0 min). Finally, fraction AAE-F16 (4.8 mg, RT 39.0 min) was separated using an isocratic mode at 25% ACN for 80 min (20 additional minutes compared to the other isocratic methods used) to yield **A8** (0.7 mg, RT 40.3 min).

### Structural elucidation of the isolated plant-derived compounds

The specific rotations of each plant-derived compounds were measured in MeOH on a JASCO P-1030 polarimeter (Loveland, CO, United States) in a 10 cm tube. NMR spectroscopic data were recorded on a Bruker Avance Neo 600 MHz NMR spectrometer equipped with a QCI 5 mm cryoprobe and a SampleJet automated sample changer (Bruker BioSpin, Rheinstetten, Germany). ^1^H-NMR experiments were recorded in DMSO-*d_6_*. Chemical shifts are reported in parts per million (δ) and coupling constants (*J*) in Hz. The residual DMSO-*d_6_* signal at δ_H_ 2.50 and δ_C_ 39.5; was used as an internal standard for ^1^H and ^13^C, respectively (*45*). HRMS acquisition parameters for isolated compounds were identical to those we previously reported (*30*). The purity of all isolated compounds was assessed by UHPLC–ELSD and UHPLC–CAD analyses.

### Cell viability assay

Cell viability of plant extract- or plant-derived compounds-treated AOAOs and A549 cells was determined by LDH release (CyQUANT LDH cytotoxicity assay, Invitrogen). Supernatants were collected and processed according to the manufacturer’s instructions, and absorbance was measured using a microplate reader (FilterMax F5 multi-mode plate reader, Molecular Devices). Viability was normalized to vehicle-treated controls.

For ALI-HAE experiments, cell viability was measured by monitoring cell metabolic activity using a resazurin-based assay as previously described (*30*, *40*). Briefly, ALI-HAE were to 3 µg/mL of resazurin (Invitrogen) for 3 h. The reduction of resazurin into fluorescent resorufin by metabolically active cells was monitored by measuring the fluorescence using a plate reader (FilterMax F5, multi-Mode Microplate Reader, Molecular Device). Cell viability was normalized with vehicle-treated controls.

## Data analysis

All calculations were performed using Excel 365 (Microsoft) and statistical analyses using Prism 10 software. All data are representative of two to four independent experiments and presented as the mean ± SEM except when mentioned in figure legends. Each test was detailed in figure legends.

## Author contributions

**Conceptualization:** Mathieu Hubert, Paola Haemmerli, Emerson Ferreira-Queiroz, Sophie Clément, Jean-Luc Wolfender and Caroline Tapparel. **Data curation:** Mathieu Hubert, Paola Haemmerli and Arnaud Gaudry. **Formal analysis:** Mathieu Hubert, Paola Haemmerli and Arnaud Gaudry. **Funding acquisition:** Jean-Luc Wolfender and Caroline Tapparel. **Investigation:** Mathieu Hubert, Paola Haemmerli, Laurence Marcourt, Emy Lara-Quintero, Loan Arthaud, Luis-Manuel Quiros-Guerrero, Sarah Donnaray and Katia Rimensberger. **Methodology:** Mathieu Hubert, Paola Haemmerli, Elodie Alessandri-Gradt, Georgios Stroulios, Yves Cambet, François Prodon and Bohumil Maco. **Project administration:** Sophie Clément, Jean-Luc Wolfender and Caroline Tapparel. **Resources:** Samuel Constant, Salvatore Simmini and Antonio Grondin. **Supervision:** Emerson Ferreira Queiroz, Sophie Clément, Jean-Luc Wolfender and Caroline Tapparel. **Validation:** Mathieu Hubert and Paola Haemmerli. **Visualization:** Mathieu Hubert, Paola Haemmerli, Luis-Manuel Quiros-Guerrero and Sarah Donnaray. **Writing – original draft:** Mathieu Hubert and Paola Haemmerli. **Writing – review & editing:** Mathieu Hubert, Paola Haemmerli, Emerson Ferreira Queiroz, Sophie Clément, Jean-Luc Wolfender and Caroline Tapparel.

## Data availability

All data are available in the main text or the Supplementary Materials. Additional data will be made available on request.

## SUPPLEMENTAL FIGURE LEGENDS

**Fig. S1.**
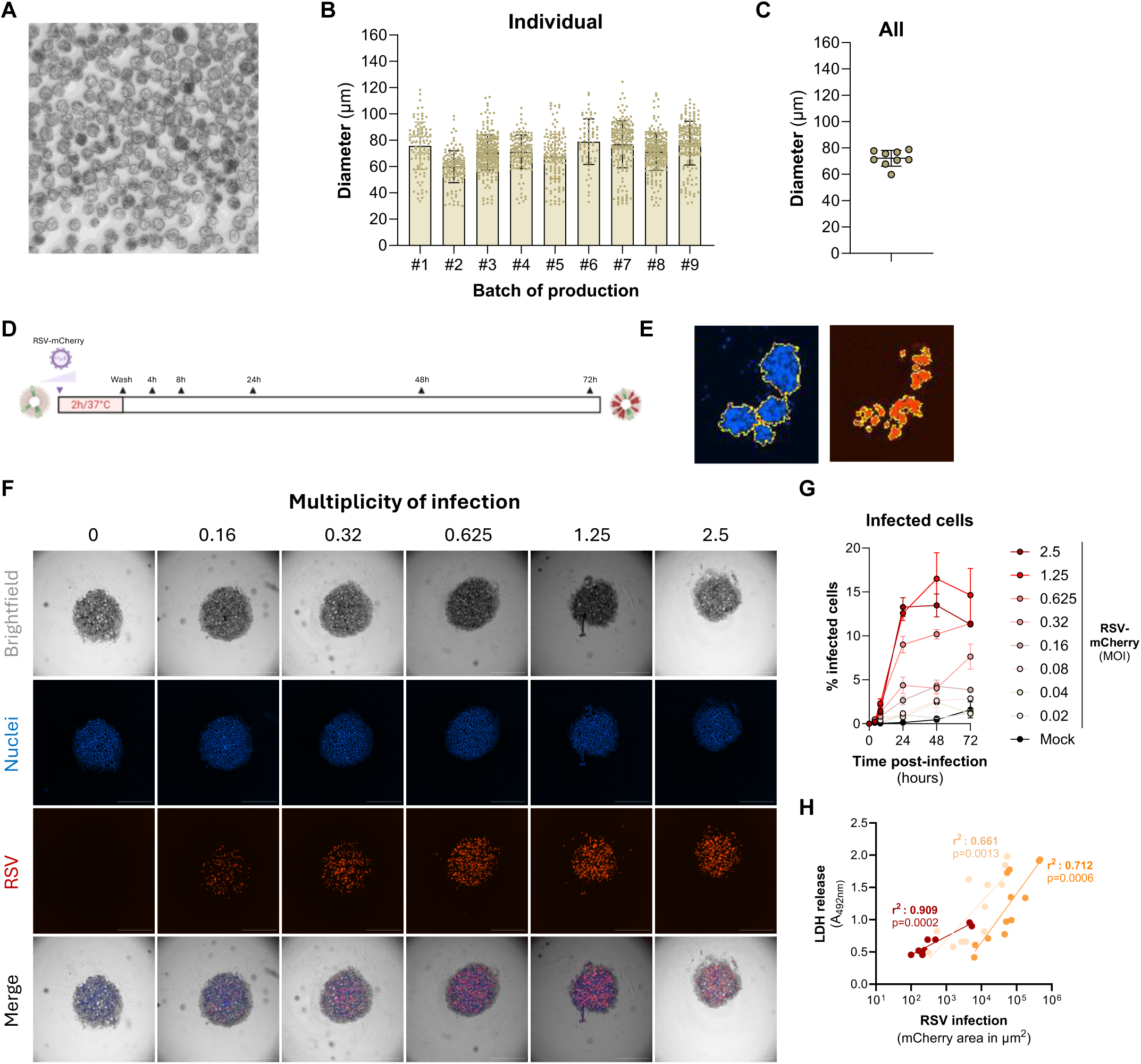
Characterization of apical-out airway organoids (AOAOs) and permissivity to RSV infection (related to Figure 1) **(A)** Brightfield image of a large population of AOAOs after 15 days of differentiation. **(B)** Nine independent productions of AOAOs were generated and size of AOAOs was measured with Cytation 5 microscope and analyzed using Gen5 software. Each dot represents a single organoid. Data are expressed as mean±SD. **(C)** Representation of the mean size of nine independent productions of AOAOs. **(D)** Schematic workflow of AOAOs infections by RSV-mCherry. Briefly, AOAOs were exposed to increasing multiplicities of infection of RSV-mCherry for 2h at 37°C. After inoculum removal, AOAOs were washed and collected at 4h, 8h, 24h, 48h and 72hpi for analysis. **(E)** Analytical pipeline for quantification of the Hoechst-positive (left) and mCherry-positive (right) areas using Gen5 software from Agilent BioTek. The yellow line shows the regions of interest. **(F)** Representative images of mock-treated and RSV-infected AOAOs acquired using a BioTek Cytation cell imaging multimode reader. Nuclei are stained in blue, RSV in red. Multiplicities of infection are indicated above images. **(G)** Flow cytometric percentage of mCherry^+^ infected cells in mock-treated and RSV-infected AOAOs. Data are expressed as mean±SEM of two independent experiments. **(H)** Correlation between mCherry area and LDH release in extracellular medium of RSV-infected AOAOs. Statistical test: simple linear regression. Data were obtained after AOAO infections with three independent viral stocks.

**Fig. S2.**
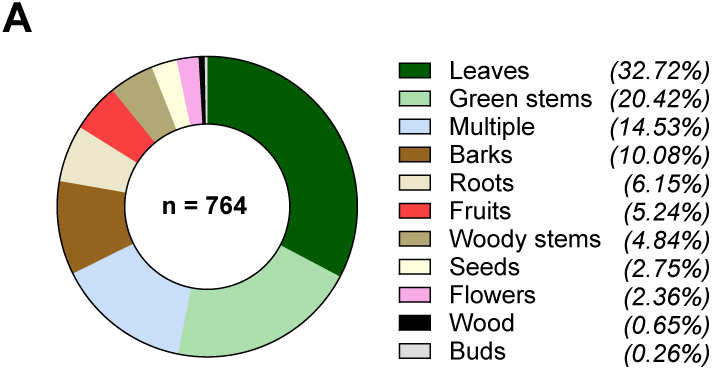
Organ diversity of the natural products (NPs) library. Repartition of organ-of-origin in percentage from which extracts were collected. n: total number of extracts.

**Fig. S3.**
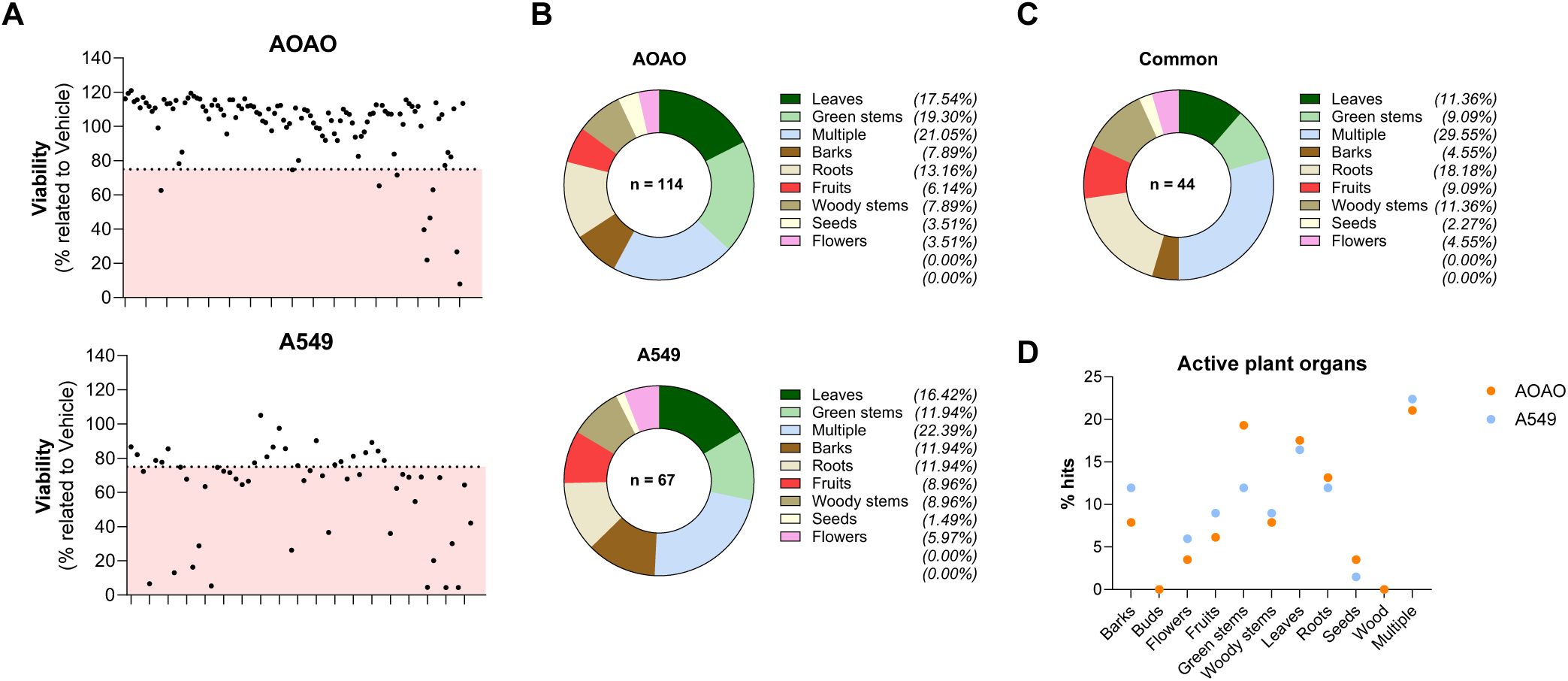
High-throughput screening of natural plant extracts against RSV in AOAOs (related to Figure 2) **(A)** Cytotoxicity of the active extracts in AOAOs (top) and A549 cells (bottom) using LDH release as a hallmark of cytotoxicity. The pink area represents the cytotoxic zone (threshold: 75%). Data were normalized to vehicle-treated controls and expressed as the mean of two independent experiments. **(B)** Repartition of AOAO-specific (top) and A549-specific (bottom) active extracts classified by plant organ. **(C)** Repartition of common active extracts in both AOAOs and A549 cells classified by plant organ. n: total number of extracts. **(D)** Distribution of AOAO- and A549-specific hits classified by plant organ. n: total number of extracts.

**Fig S4.**
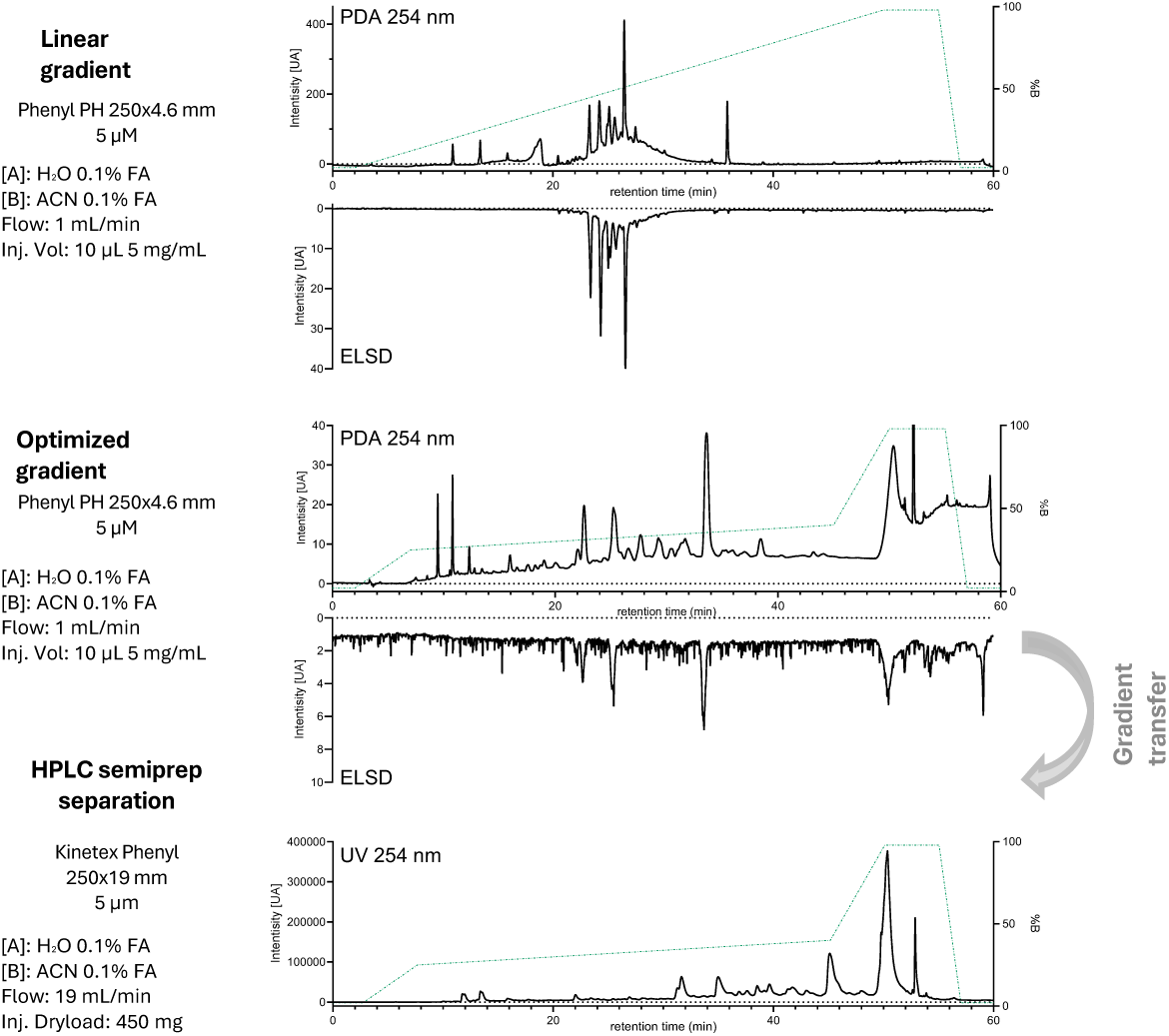
Chromatographic optimization and analytical-to-semi-preparative transfer for the isolation of compounds from the *Ampelocissus arachnoida* ethyl acetate extract. **(A)** HPLC–PDA–ELSD chromatograms of the *Ampelocissus arachnoida* ethyl acetate extract acquired with an initial linear gradient (top), an optimized gradient (middle), and after transfer of the optimized conditions to the semi-preparative flash scale (bottom). **(B)** Inverted chromatographic traces correspond to the ELSD detector.

**Fig S5.**
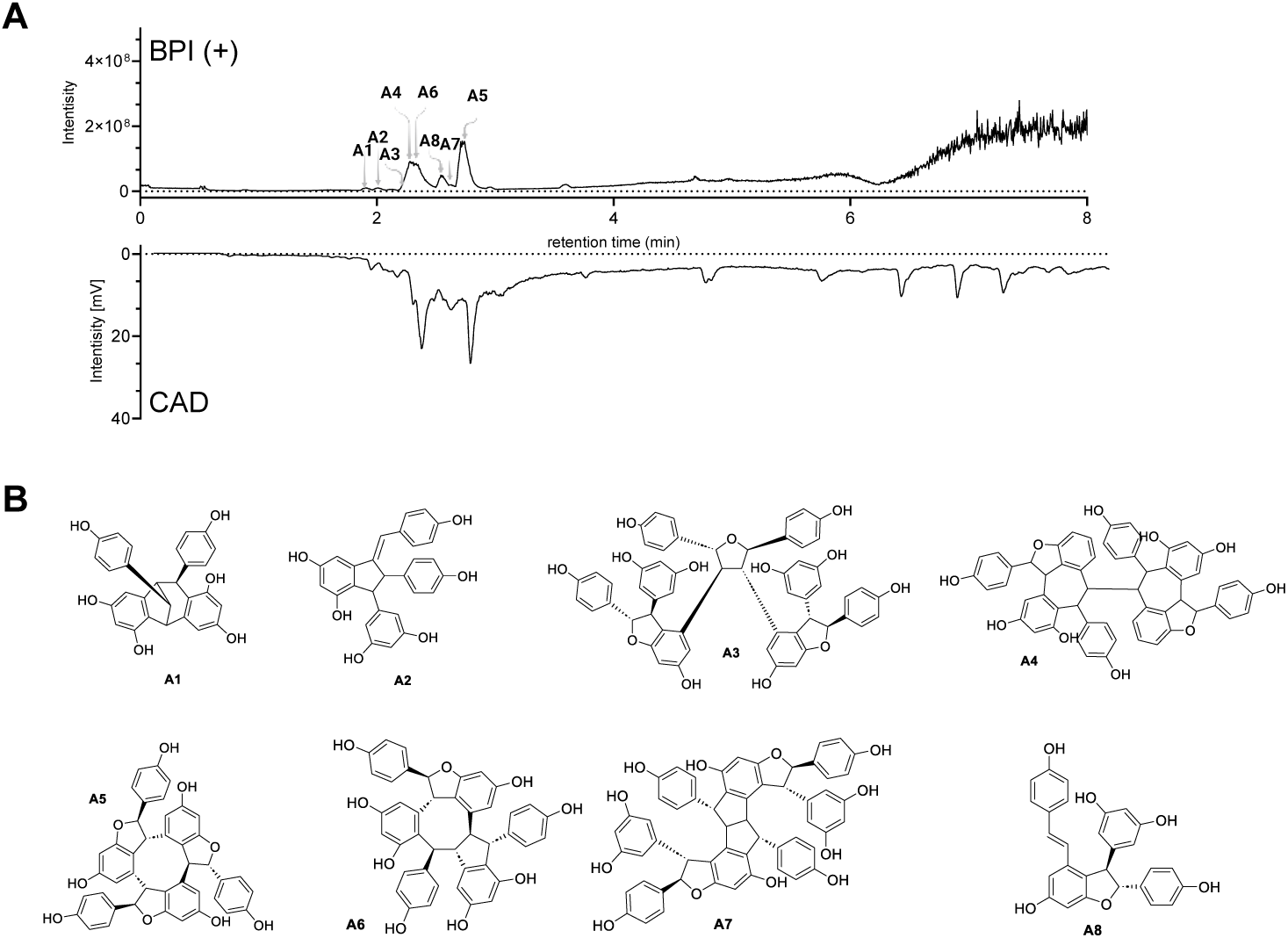
UHPLC–HRMS/CAD profiling and compound isolation from the *Ampelocissus arachnoida* ethyl acetate extract. **(A)** Base peak intensity (BPI) chromatogram acquired in positive ionization mode by UHPLC–HRMS, overlaid with the semi-quantitative charged aerosol detector (CAD) trace, recorded over an 8-min run on a Waters BEH C18 column (150 × 2.1 mm, 1.7 μm) using a linear gradient from 5 to 100% acetonitrile. Numbers above peaks correspond to isolated compounds. **(B)** Chemical structures of the isolated compounds (**A1–A8**) obtained from the *Ampelocissus arachnoida* ethyl acetate extract.

**Fig S6.**
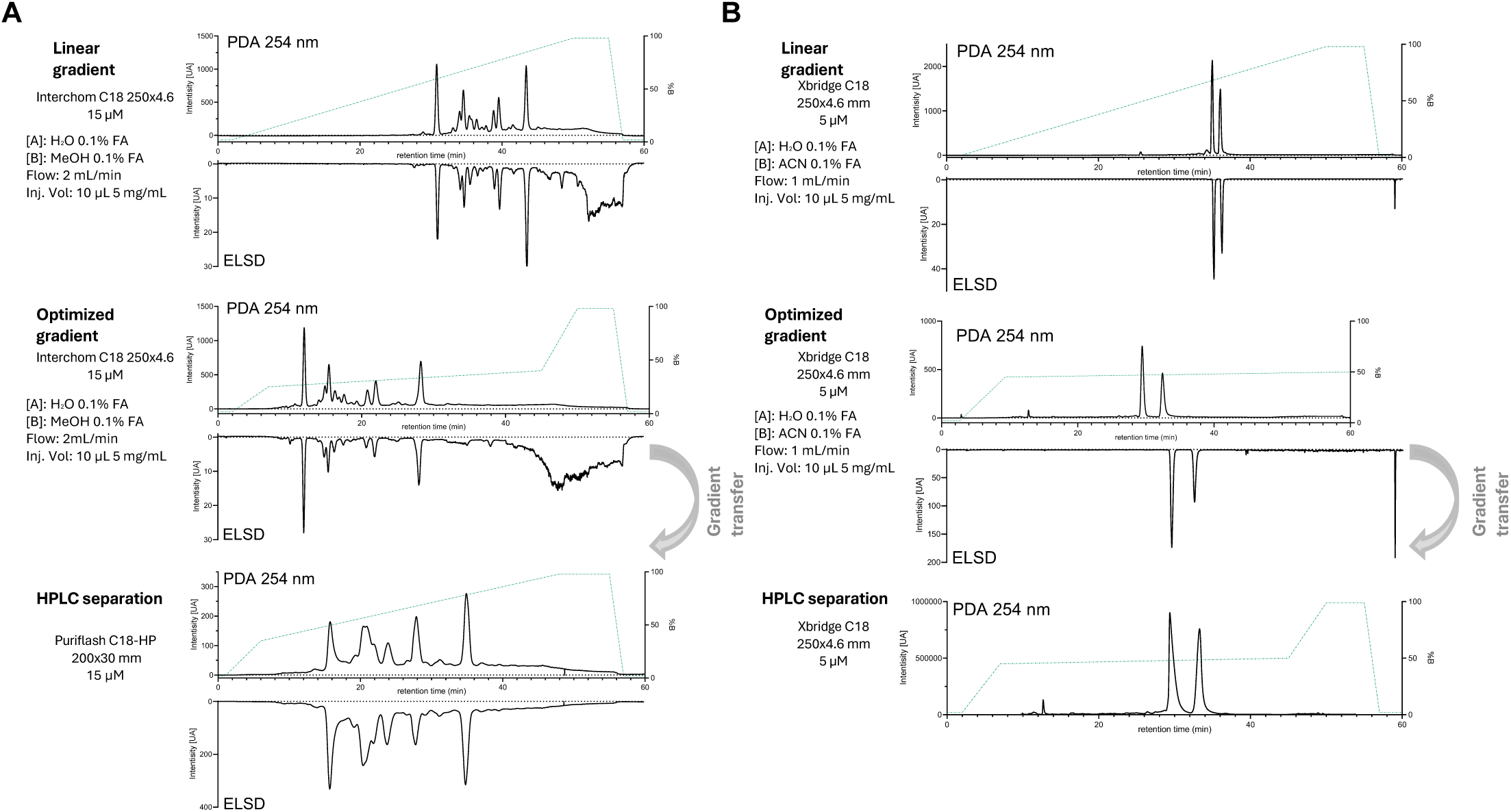
Chromatographic optimization and scale-up for the isolation of compounds from the *Clausena wallichii* ethyl acetate extract. **(A)** HPLC–PDA–ELSD chromatograms of the *Clausena wallichii* ethyl acetate extract recorded with an initial linear gradient (top), an optimized gradient (middle), and after transfer of the optimized conditions to the semi-preparative flash scale (bottom). **(B)** HPLC–PDA–ELSD chromatograms of fraction *Clausena wallichii* ethyl acetate-F29 acquired under the same conditions: linear gradient (top), optimized gradient (middle), and semi-preparative flash scale (bottom). Inverted chromatographic traces correspond to the ELSD detector.

**Fig S7.**
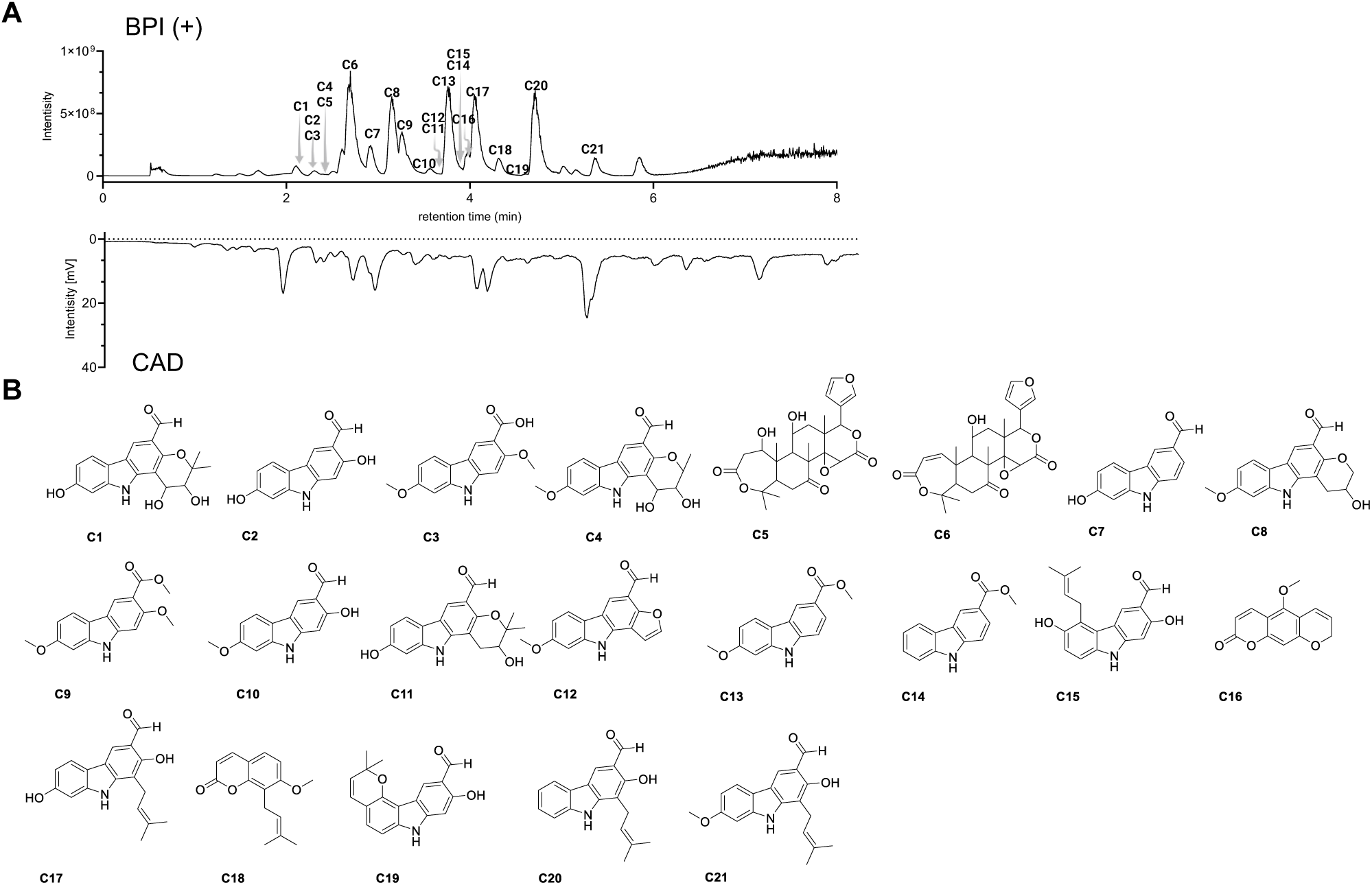
UHPLC–HRMS/CAD profiling and compound isolation from the *Clausena wallichii* ethyl acetate extract. **(A)** Base peak intensity (BPI) chromatogram acquired in positive ionization mode by UHPLC–HRMS, overlaid with the semi-quantitative charged aerosol detector (CAD) trace, recorded over an 8-min run on a Waters BEH C18 column (150 × 2.1 mm, 1.7 μm) using a linear gradient from 5 to 100% acetonitrile. Numbers above peaks correspond to isolated compounds. **(B)** Chemical structures of the isolated compounds (**C1–C21**) obtained from the *Clausena wallichii* ethyl acetate extract.

**Fig S8.**
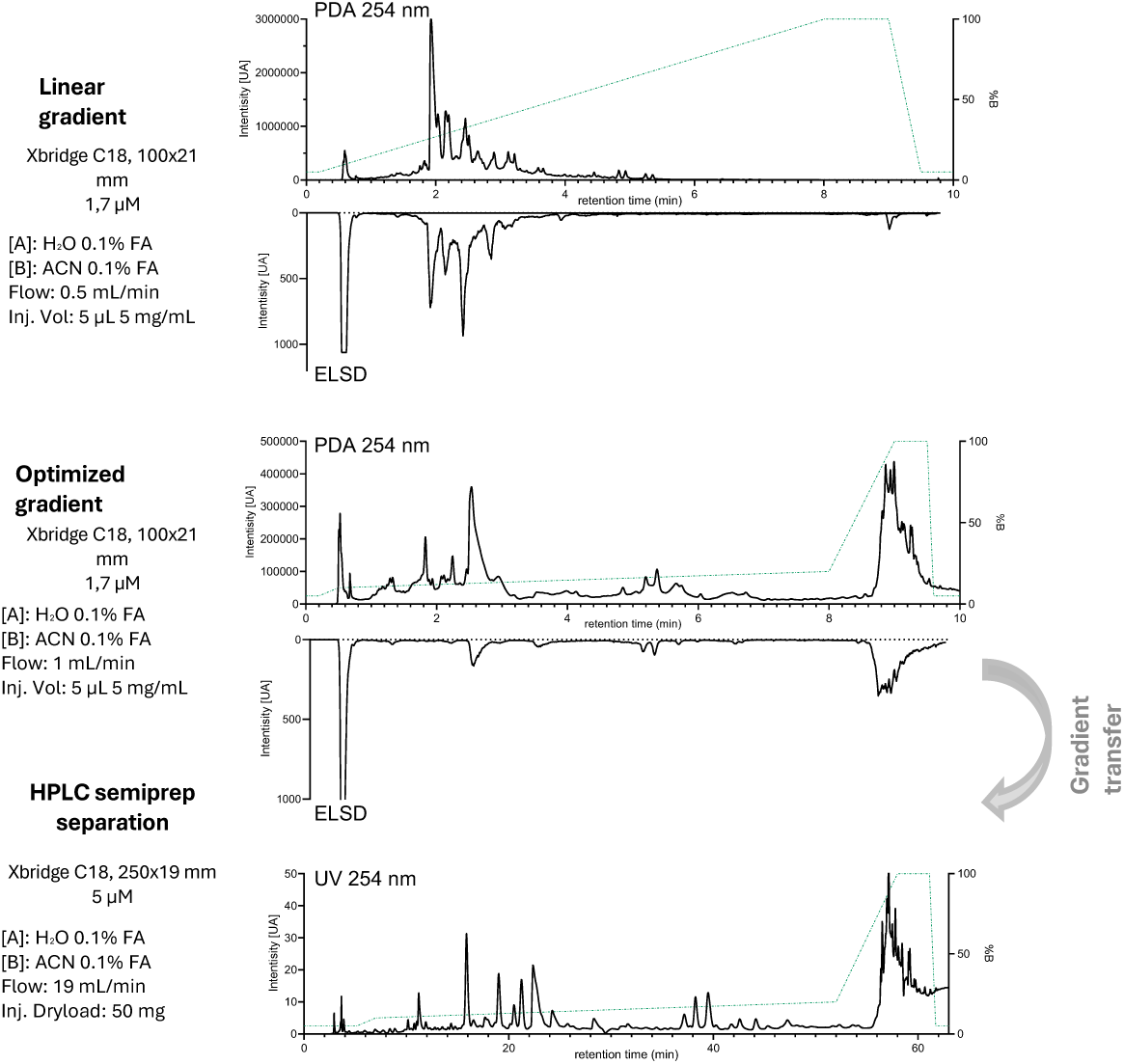
Chromatographic optimization and analytical-to-semi-preparative transfer for the isolation of compounds from the *Litsea polyantha* ethyl acetate extract. HPLC–PDA–ELSD chromatograms of the *Litsea polyantha*ethyl acetate extract acquired with an initial linear gradient (top), an optimized gradient (middle), and after transfer of the optimized conditions to the semi-preparative scale (bottom). Inverted chromatographic traces correspond to the ELSD detector.

**Fig S9.**
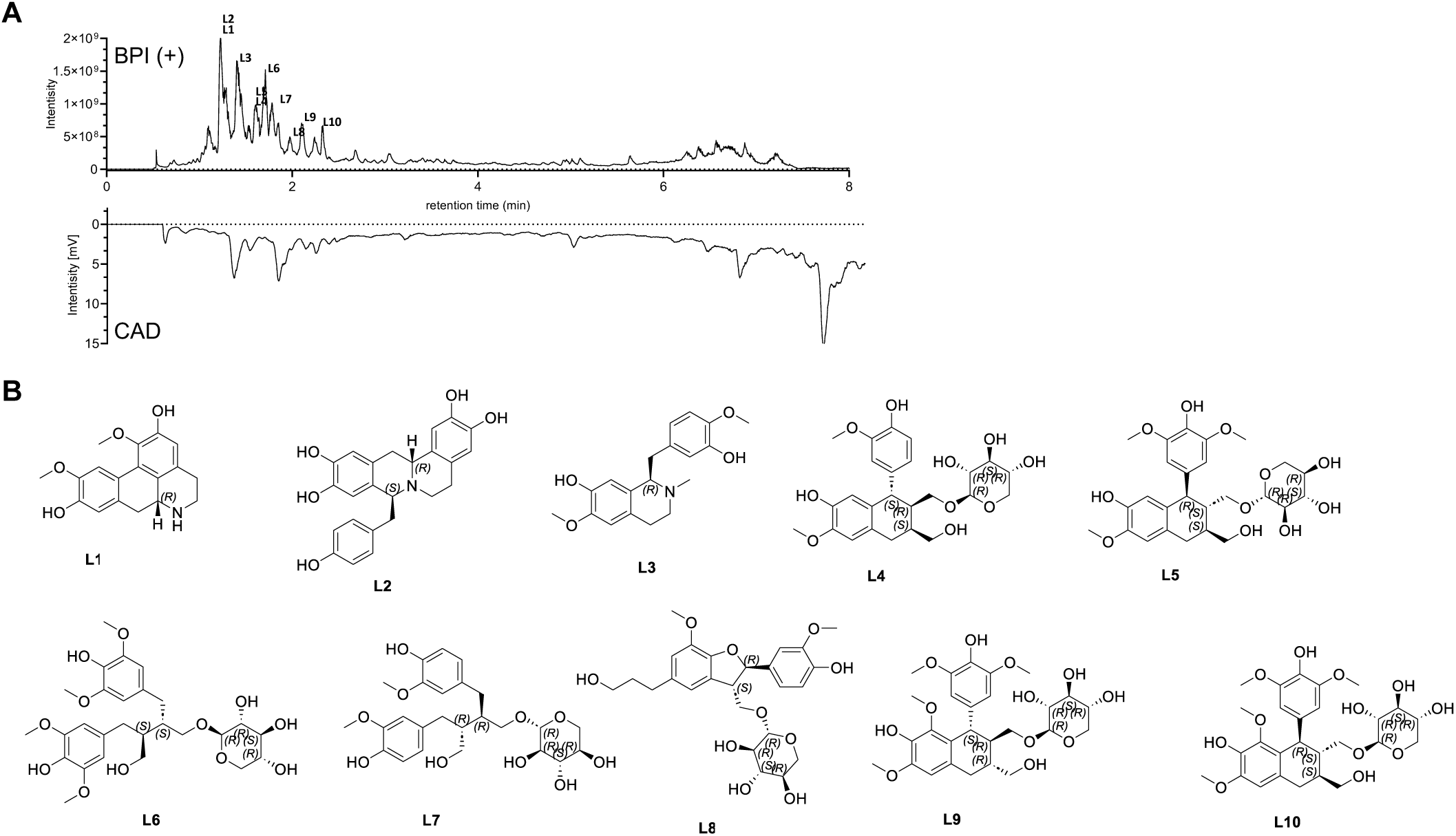
UHPLC–HRMS/CAD profiling and compound isolation from the *Litsea polyantha* ethyl acetate extract. **(A)** Base peak intensity (BPI) chromatogram acquired in positive ionization mode by UHPLC–HRMS, overlaid with the semi-quantitative charged aerosol detector (CAD) trace, recorded over an 8-min run on a Waters BEH C18 column (150 × 2.1 mm, 1.7 μm) using a linear gradient from 5 to 100% acetonitrile. Numbers above peaks correspond to isolated compounds. **(B)** Chemical structures of the isolated compounds (L1–L10) obtained from the *Litsea polyantha* ethyl acetate extract.

**Fig S10.**
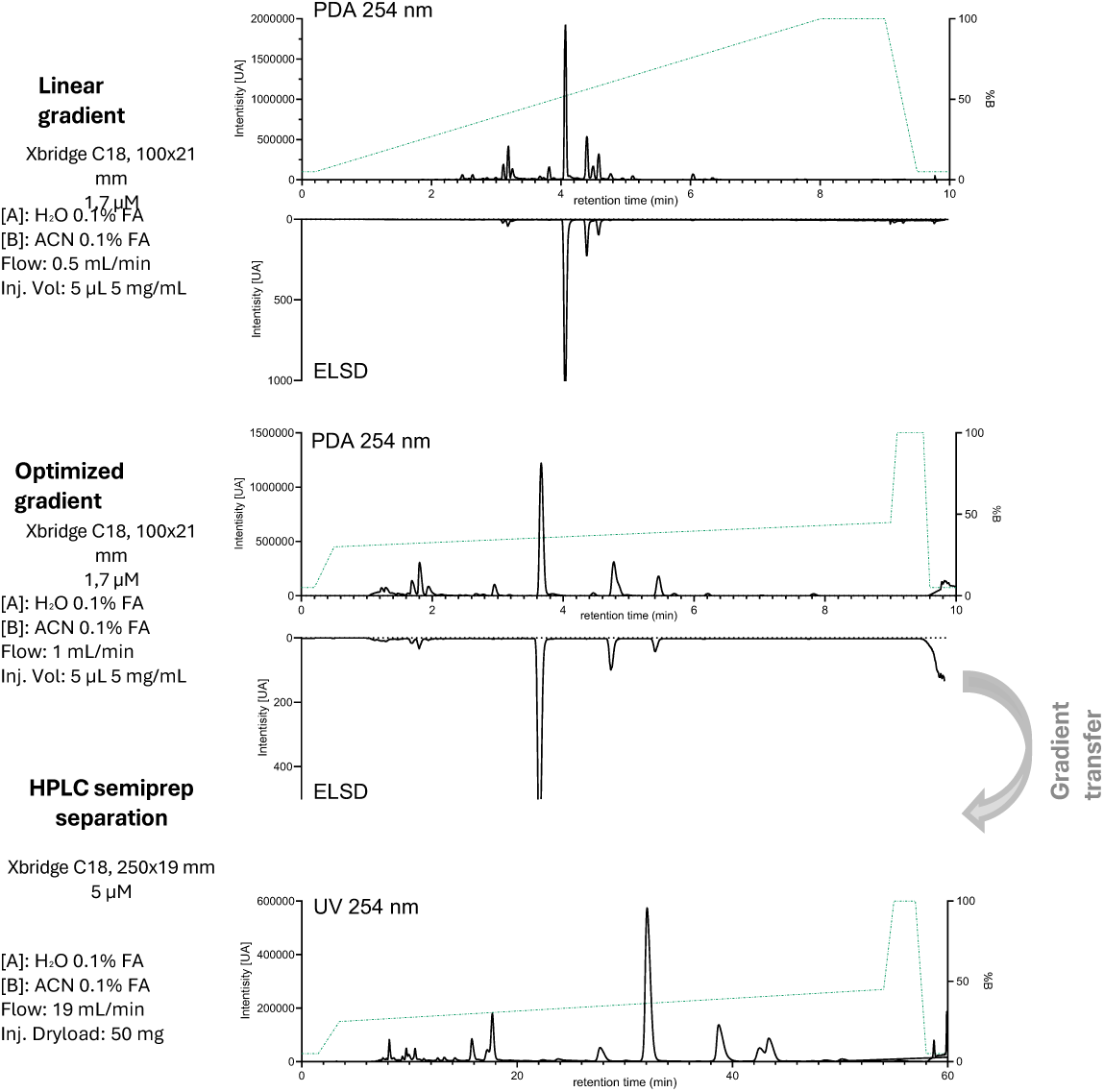
Chromatographic optimization and analytical-to-semi-preparative transfer for the isolation of compounds from the *Vepris macrophylla* ethyl acetate extract. HPLC–PDA–ELSD chromatograms of the *Vepris macrophylla* ethyl acetate extract acquired with an initial linear gradient (top), an optimized gradient (middle), and after transfer of the optimized conditions to the semi-preparative scale (bottom). Inverted chromatographic traces correspond to the ELSD detector.

**Fig S11.**
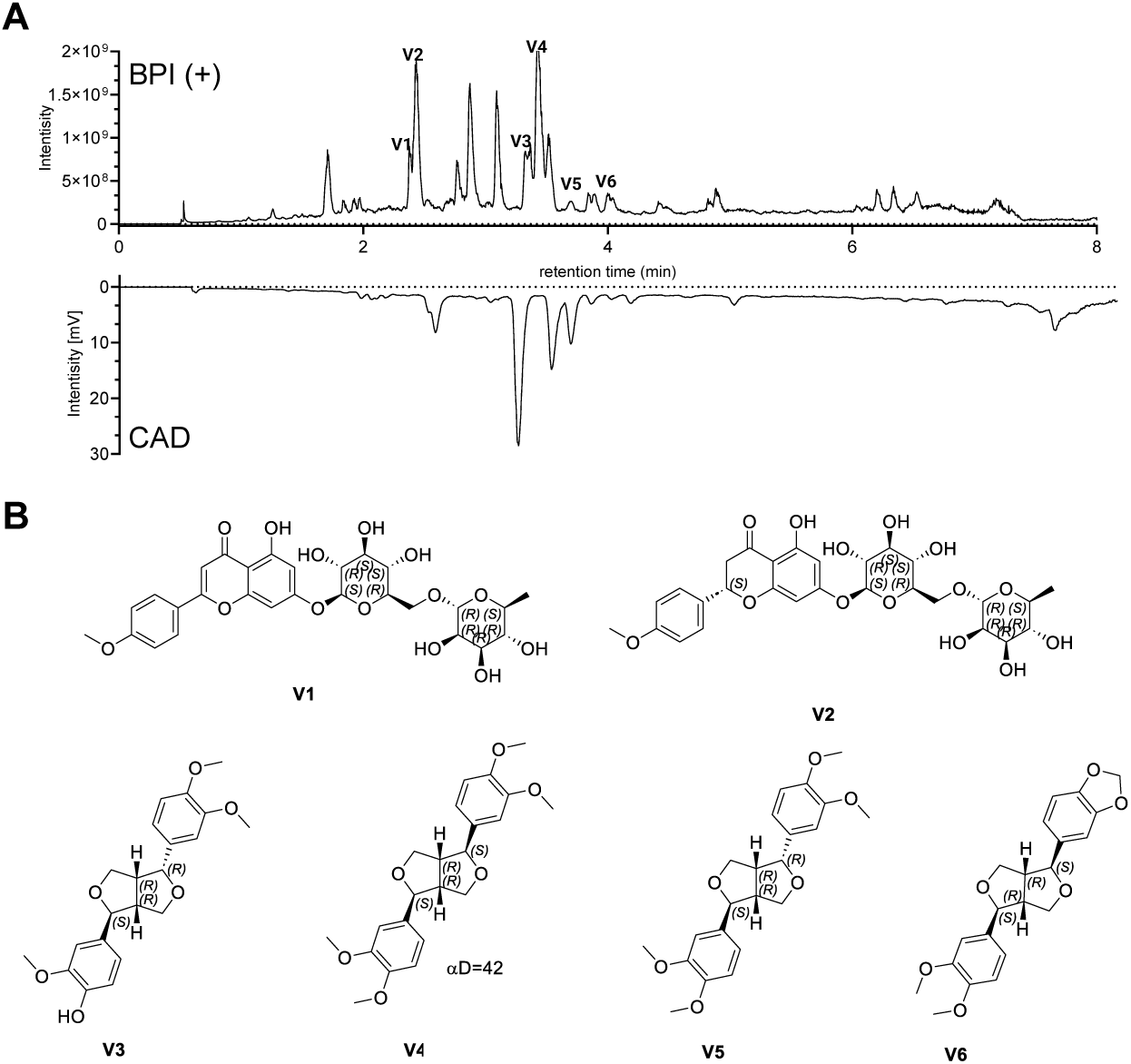
UHPLC–HRMS/CAD profiling and compound isolation from the *Vepris macrophylla* ethyl acetate extract. **(A)** Base peak intensity (BPI) chromatogram acquired in positive ionization mode by UHPLC–HRMS, overlaid with the semi-quantitative charged aerosol detector (CAD) trace, recorded over an 8-min run on a Waters BEH C18 column (150 × 2.1 mm, 1.7 μm) using a linear gradient from 5 to 100% acetonitrile. Numbers above peaks correspond to isolated compounds. **(B)** Chemical structures of the isolated compounds (**V1–V6**) obtained from the *Vepris macrophylla* ethyl acetate extract.

**Fig. S12.**
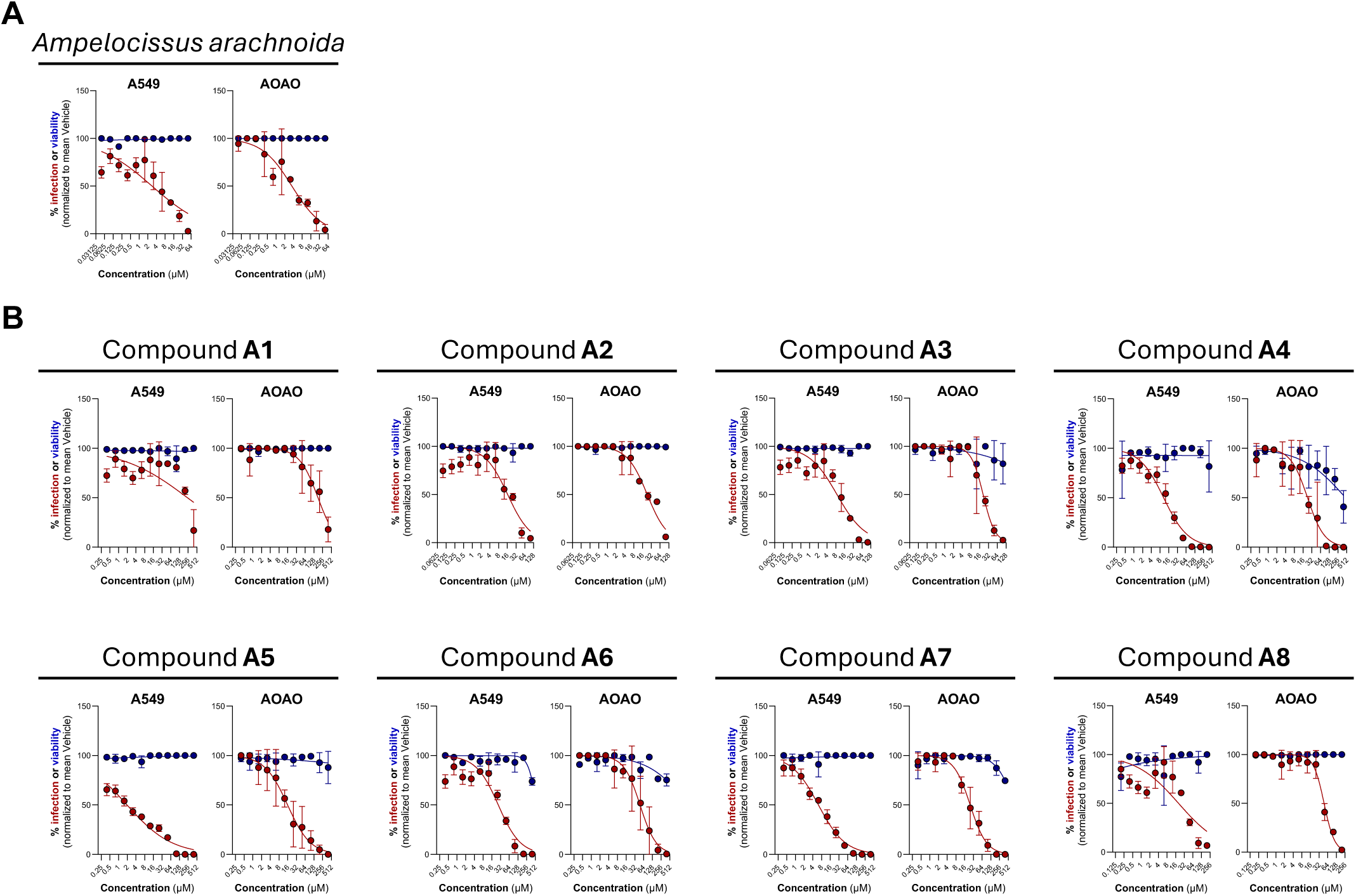
Cytotoxicity and antiviral activity of *Ampelocissus arachnoida*-derived compounds against RSV. Dose-response analysis of cytotoxicity (blue curve) and infection (red curve) of A549 cells (left) and AOAOs (right) in the presence of increasing doses of the *Ampelocissus arachnoida* ethyl acetate extract **(A)** and *Ampelocissus arachnoida*-derived compounds **(B)**. 50% cytotoxic (CC₅₀) and inhibitory (IC₅₀) concentrations are reported in Table S2.

**Fig. S13.**
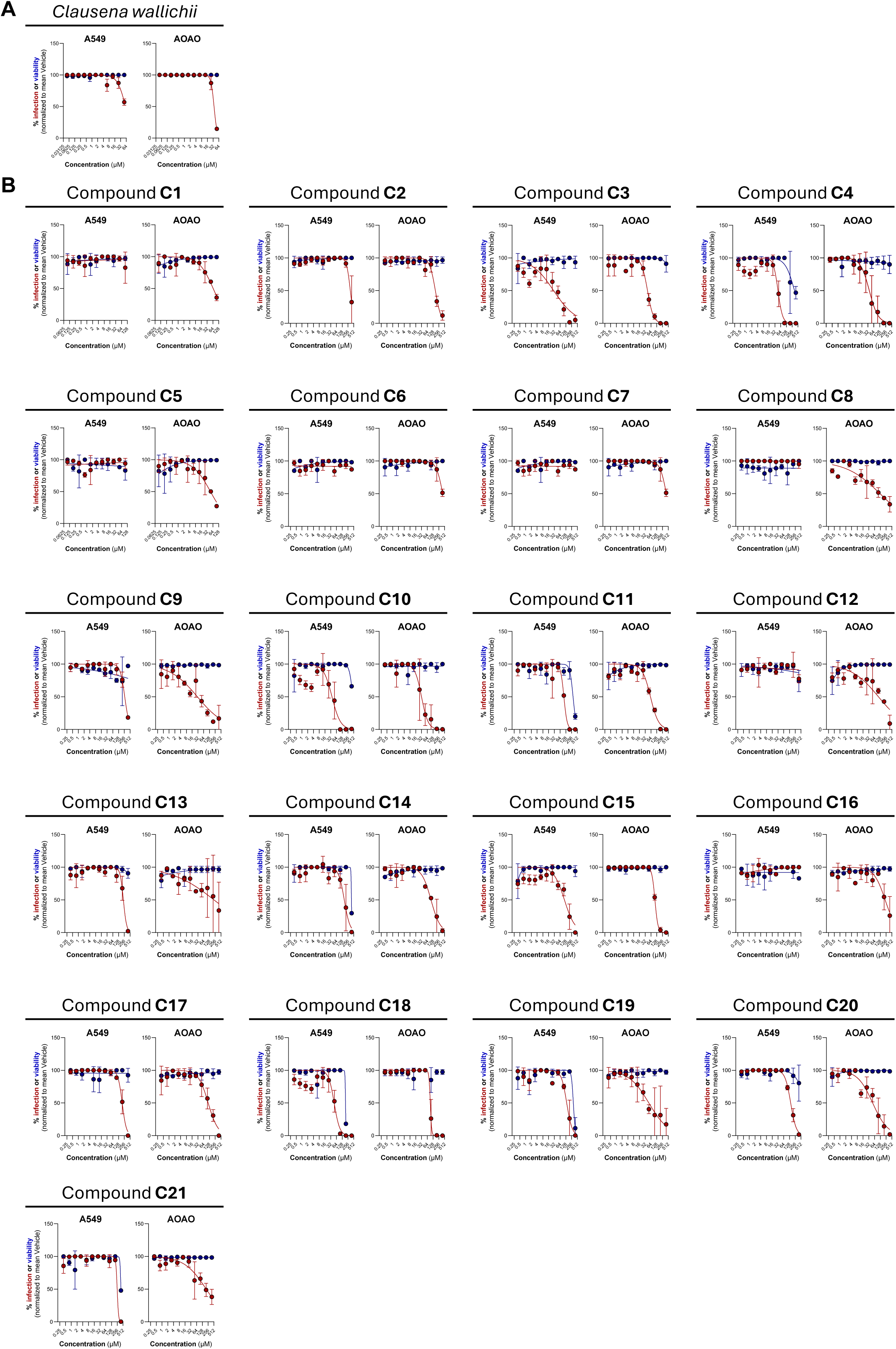
Antiviral activity of *Clausena wallichii*-derived compounds against RSV. Dose-response analysis of cytotoxicity (blue curve) and infection (red curve) of A549 cells (left) and AOAOs (right) in the presence of increasing doses of the *Clausena wallichii* ethyl acetate extract **(A)** and *Clausena wallichii*-derived compounds **(B)**. 50% cytotoxic (CC₅₀) and inhibitory (IC₅₀) concentrations are reported in Table S2.

**Fig. S14.**
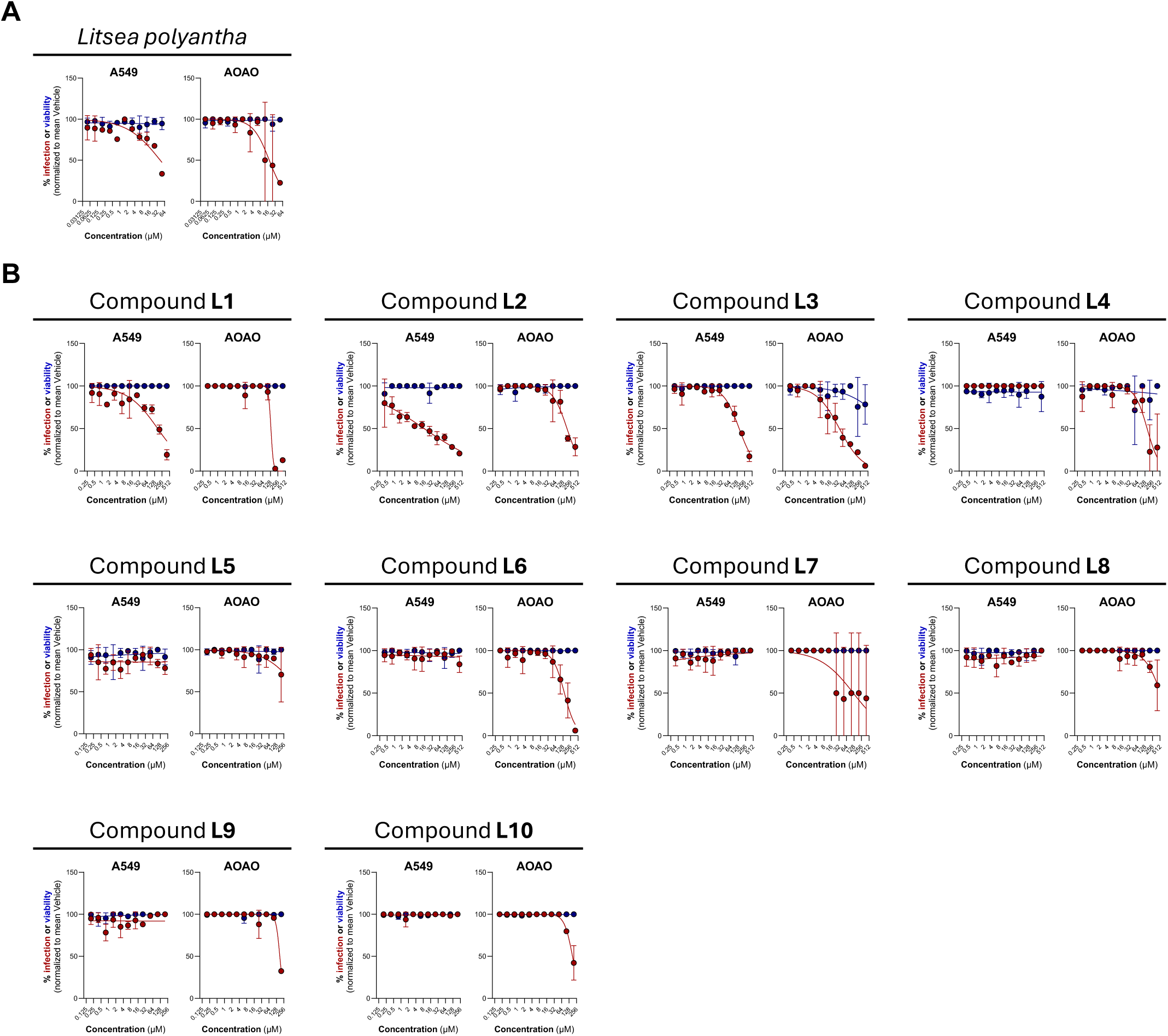
Cytotoxicity and antiviral activity of *Litsea polyantha*-derived compounds against RSV. Dose-response analysis of cytotoxicity (blue curve) and infection (red curve) of A549 cells (left) and AOAOs (right) in the presence of increasing doses of the *Litsea polyantha* ethyl acetate extract **(A)** and *Litsea polyantha*-derived compounds **(B)**. 50% cytotoxic (CC₅₀) and inhibitory (IC₅₀) concentrations are reported in Table S2.

**Fig. S15.**
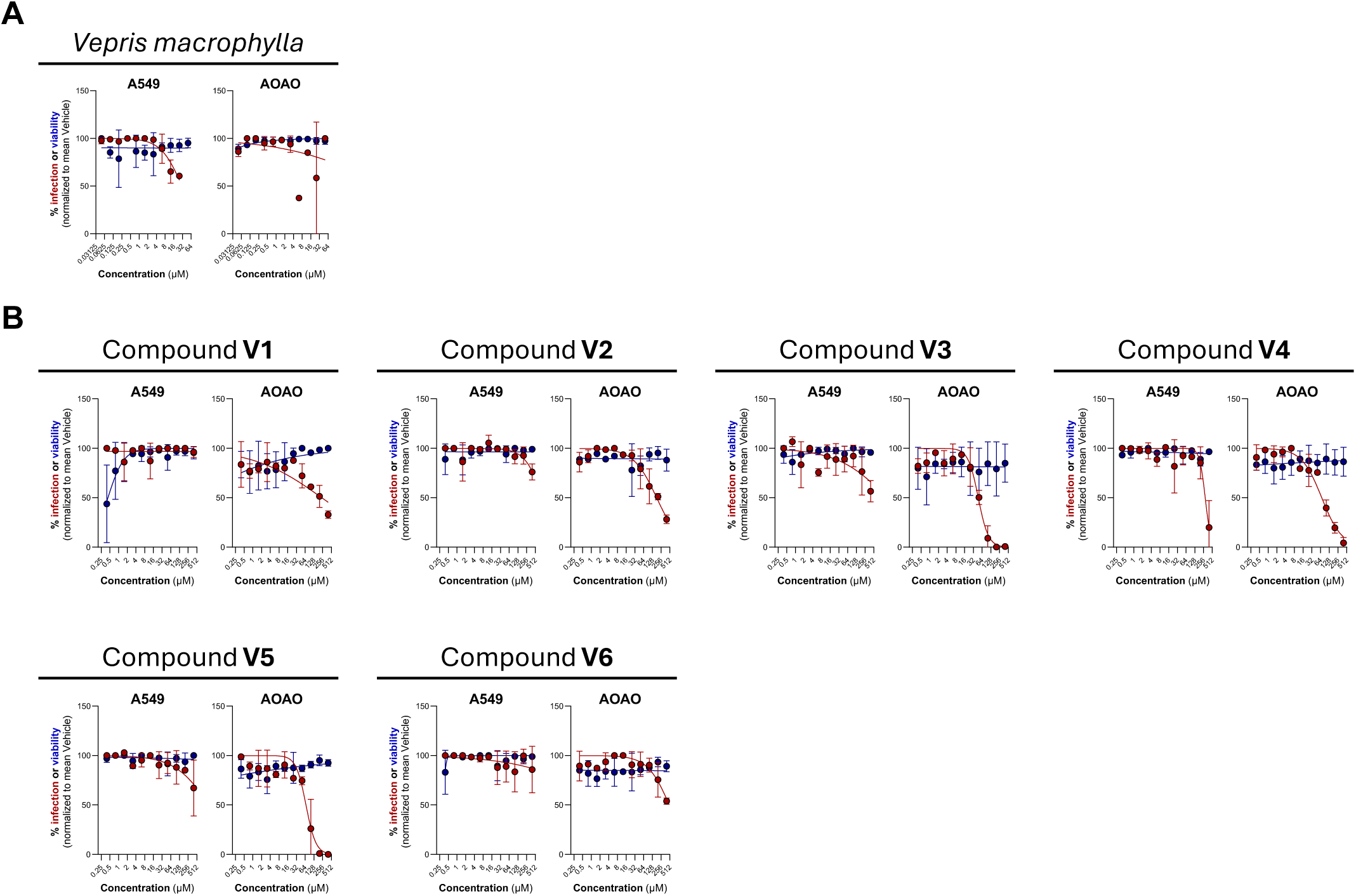
Cytotoxicity and antiviral activity of *Vepris macrophylla*-derived compounds against RSV. Dose-response analysis of cytotoxicity (blue curve) and infection (red curve) of A549 cells (left) and AOAOs (right) in the presence of increasing doses of the *Vepris macrophylla* ethyl acetate extract **(A)** and *Vepris macrophylla*-derived compounds **(B)**. 50% cytotoxic (CC₅₀) and inhibitory (IC₅₀) concentrations are reported in Table S2.

**Fig S16.**
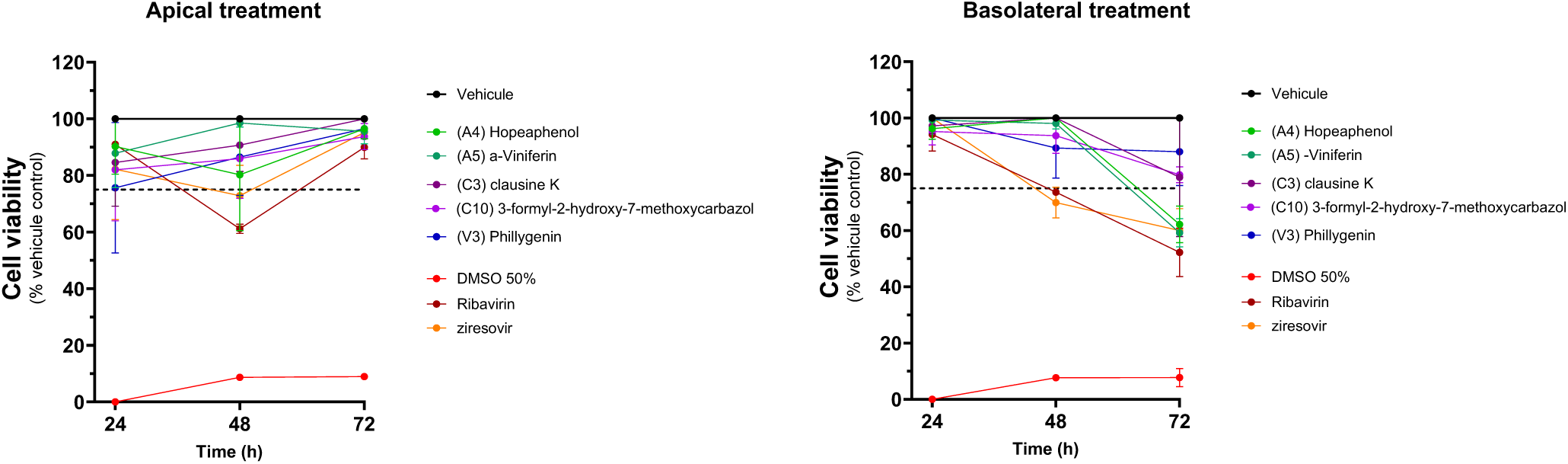
Cell viability of human airway epithelia reconstituted at the air-liquid interface (ALI-HAE) and treated with plant-derived compounds (Related to Figure 4) Air-liquid interface-cultures human airway epithelia (ALI-HAE) were daily exposed to the maximal antiviral but non-cytotoxic concentration of each compound via the apical (left) or basolateral (right) compartment. Cytotoxicity was assessed using a resazurin-based cell viability assay by quantifying resorufin production in the apical supernatant. Data are expressed as mean ± SD from two independent experiments.

**Table S1.**
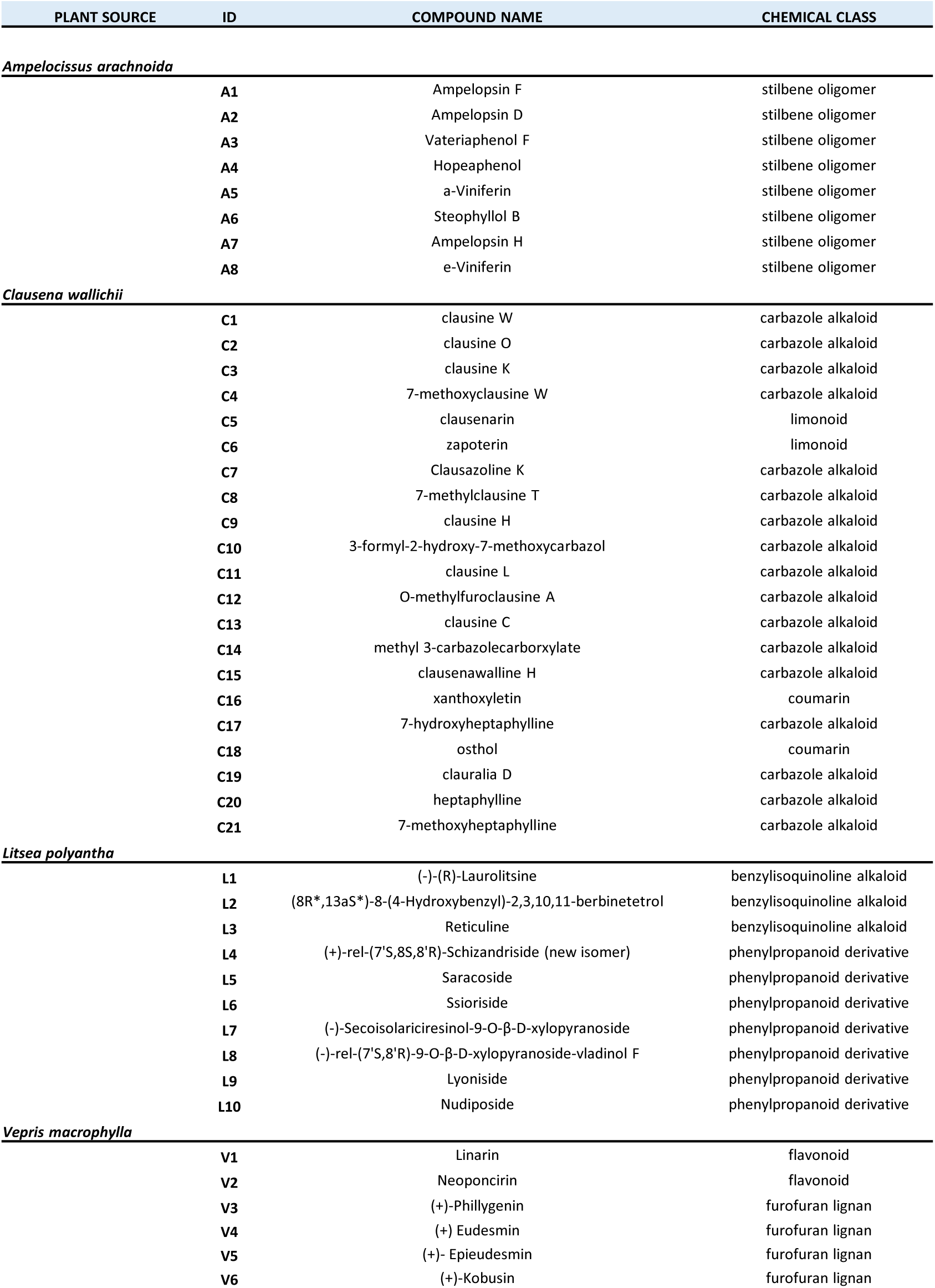
Characteristics of plant-derived isolated compounds. Plant-derived isolated compounds classified by plant of origine, name and chemical class.

| PLANT SOURCE | ID | COMPOUND NAME | CHEMICAL CLASS |
| --- | --- | --- | --- |
| *Ampelocissus arachnoida* |  |  |  |
|  | A1 | Ampelopsin F | stilbene oligomer |
|  | A2 | Ampelopsin D | stilbene oligomer |
|  | A3 | Vateriaphenol F | stilbene oligomer |
|  | A4 | Hopeaphenol | stilbene oligomer |
|  | A5 | a-Viniferin | stilbene oligomer |
|  | A6 | Steophyllol B | stilbene oligomer |
|  | A7 | Ampelopsin H | stilbene oligomer |
|  | A8 | e-Viniferin | stilbene oligomer |
| *Clausena wallichii* |  |  |  |
|  | C1 | clausine W | carbazole alkaloid |
|  | C2 | clausine O | carbazole alkaloid |
|  | C3 | clausine K | carbazole alkaloid |
|  | C4 | 7-methoxyclausine W | carbazole alkaloid |
|  | C5 | clausenarin | limonoid |
|  | C6 | zapoterin | limonoid |
|  | C7 | Clausazoline K | carbazole alkaloid |
|  | C8 | 7-methylclausine T | carbazole alkaloid |
|  | C9 | clausine H | carbazole alkaloid |
|  | C10 | 3-formyl-2-hydroxy-7-methoxycarbazol | carbazole alkaloid |
|  | C11 | clausine L | carbazole alkaloid |
|  | C12 | O-methylfuroclausine A | carbazole alkaloid |
|  | C13 | clausine C | carbazole alkaloid |
|  | C14 | methyl 3-carbazolecarboxylate | carbazole alkaloid |
|  | C15 | clausenawalline H | carbazole alkaloid |
|  | C16 | xanthoxyletin | coumarin |
|  | C17 | 7-hydroxyheptaphylline | carbazole alkaloid |
|  | C18 | osthol | coumarin |
|  | C19 | clauralia D | carbazole alkaloid |
|  | C20 | heptaphylline | carbazole alkaloid |
|  | C21 | 7-methoxyheptaphylline | carbazole alkaloid |
| *Litsea polyantha* |  |  |  |
|  | L1 | (-)-(R)-Laurolitsine | benzylisoquinoline alkaloid |
|  | L2 | (8R*,13aS*)-8-(4-Hydroxybenzyl)-2,3,10,11-berbinetrol | benzylisoquinoline alkaloid |
|  | L3 | Reticuline | benzylisoquinoline alkaloid |
|  | L4 | (+)-rel-(7'S,8S,8'R)-Schizandriside (new isomer) | phenylpropanoid derivative |
|  | L5 | Saracoside | phenylpropanoid derivative |
|  | L6 | Ssioriside | phenylpropanoid derivative |
|  | L7 | (-)-Secoisolariciresinol-9-O-β-D-xylopyranoside | phenylpropanoid derivative |
|  | L8 | (-)-rel-(7'S,8'R)-9-O-β-D-xylopyranoside-vladinol F | phenylpropanoid derivative |
|  | L9 | Lyoniside | phenylpropanoid derivative |
|  | L10 | Nudiposide | phenylpropanoid derivative |
| *Vepris macrophylla* |  |  |  |
|  | V1 | Linarin | flavonoid |
|  | V2 | Neoponcirin | flavonoid |
|  | V3 | (+)-Phillygenin | furofuran lignan |
|  | V4 | (+) Eudesmin | furofuran lignan |
|  | V5 | (+)- Epieudesmin | furofuran lignan |
|  | V6 | (+)-Kobusin | furofuran lignan |

**Table S2.**
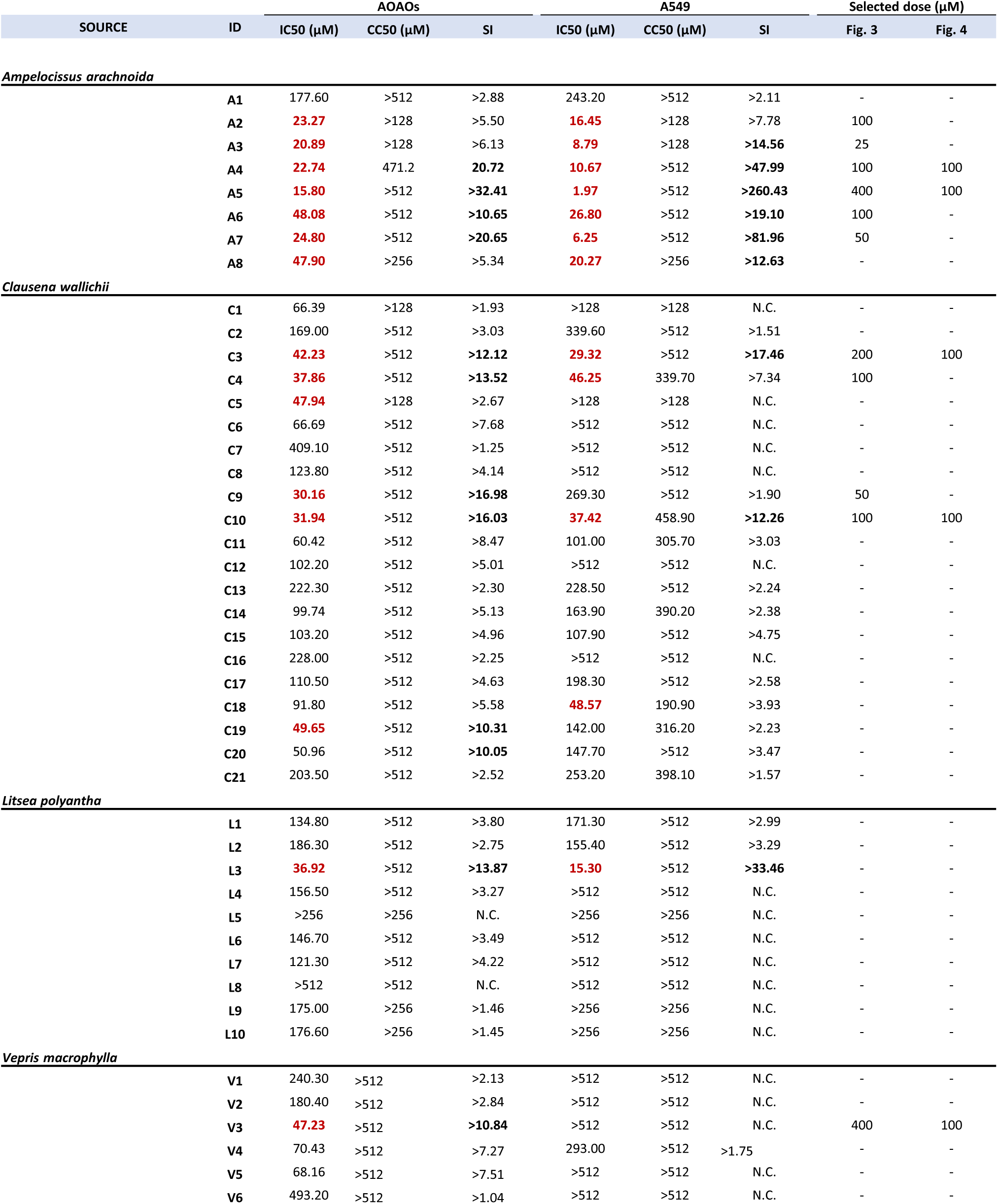
Summary of antiviral activity of plant-derived isolated compounds. Plant-derived isolated compounds antiviral activity expressed as the inhibitory concentration 50% (IC_50_), cytotoxic concentration 50% (CC_50_) and selectivity index (SI) in AOAOs and A549 cells. IC_50_ values <50μM are shown in red. Selectivity indexes >10 are shown in bold. Data represents the mean of two independent experiments. N.C.: not calculable.

